# Value-guided attention links what we learn to how much we learn

**DOI:** 10.64898/2026.08.25.747046

**Authors:** Atlas Shahamati, Alireza Soltani

**Affiliations:** Department of psychological and brain sciences, Dartmouth College, Hanover, NH, USA

**Author notes:** Corresponding author: AS, Department of Psychological and Brain Sciences, Dartmouth College, Hanover NH 03755.

**Keywords:** credit assignment, learning rate, saliency, eye-tracking, behavioral manipulation

## Abstract

Learning in uncertain environments requires identifying the relevant associations between stimuli, actions, and outcomes and determining how strongly to update these associations. Although often treated separately, these components likely interact in the brain. We hypothesized that this interaction shapes individual learning rates according to cue-choice alignment and reward outcome, thereby improving discrimination between competing cues. We tested this hypothesis using a probabilistic learning task in which human participants predicted outcomes based on multiple cues and reward feedback. We measured gaze and manipulated cue saliency to assess and influence which cues were preferentially processed during choice and feedback. Computational modeling revealed that learning rates were selectively enhanced for cues supporting the chosen option after reward and for cues opposing it after no reward. This learning-rate asymmetry based on cue-choice alignment sharpened discrimination among predictive cues, increased robustness to noise, and improved performance. Moreover, differential gaze toward supporting and opposing cues predicted this asymmetry, which was causally altered by manipulating cue saliency. Together, our results suggest that attention provides a unifying mechanism for coordinating what we learn from with how much we learn, helping preserve distinctions among competing cues and bringing several learning asymmetries within a common framework.

## Introduction

Learning in uncertain, naturalistic environments poses two main challenges: identifying the relevant associations between stimuli, actions, and reward outcomes, and determining how strongly those associations should be updated in response to new evidence and outcomes. These correspond to the problems of credit assignment (what caused the outcome) and adaptive learning-rate control (how much to update), respectively (Behrens et al., 2007; Nassar et al., 2012; Payzan-LeNestour et al., 2013; Schultz et al., 1997; Soltani et al., 2021; Soltani & Koechlin, 2021; Sutton & Barto, 1998). In reinforcement learning (RL), credit assignment and learning-rate control are often formalized as distinct components of learning, with the former determining which stimuli or actions are updated and the latter controlling the magnitude of those updates. Although they can interact within specific RL algorithms, this separation allows models to distinguish the structure of credit assignment from update dynamics (Sutton & Barto, 1998). In biological learning, however, the processes that determine what receives credit may themselves influence how strongly it is updated.

This could arise for several reasons. At the cognitive level, multiple processes become engaged as learning unfolds to resolve the credit assignment problem, and each can in turn shape learning rates (Collins & Frank, 2012; Farashahi, Rowe, et al., 2017; Moran et al., 2021; Niv et al., 2015; Yoo & Collins, 2022). Notably, despite variability across experimental conditions and reward environments (Behrens et al., 2007; Donahue & Lee, 2015; Farashahi et al., 2019; McGuire et al., 2014; Nassar et al., 2012; Simoens et al., 2024), recent studies have identified systematic patterns in learning rates driven by reward history, choice, expectations, beliefs, and outcomes, suggesting that the assignment of credit influences how strongly learning proceeds. For example, learning rates are typically higher for rewarded than unrewarded outcomes of the chosen option, with the opposite pattern for the unchosen option, a choice-confirmation bias that favors information supporting one’s choices (Lefebvre et al., 2022; Palminteri et al., 2017). Similarly, when cues conveyed others’ beliefs, learning rates were higher for prediction errors consistent with those beliefs and lower for conflicting errors, revealing confirmation bias in expectation-based learning (Yazdanpanah et al., 2026). Although it remains unclear whether high-level confirmation biases share neural mechanisms with choice-confirmation bias (Palminteri & Lebreton, 2022), learning rates are clearly shaped by multiple cognitive processes, including working memory (Collins & Frank, 2012; Yoo & Collins, 2022), beliefs (Palminteri et al., 2017; Yazdanpanah et al., 2026) and attention (Leong et al., 2017; Niv et al., 2015; Wang & Soltani, 2025).

At the synaptic level, RL is thought to rely on reward-dependent synaptic plasticity (Izhikevich, 2007; Legenstein et al., 2008; J. N. J. Reynolds & Wickens, 2002; Schultz, 2013; Soltani & Wang, 2006, 2008). Because this plasticity also depends on neural activity, learning is influenced by how strongly cues are represented during decision making and when reward feedback is received (Izhikevich, 2007; Legenstein et al., 2008; Soltani & Wang, 2006, 2008). For example, imagine deciding whether to turn left or right while trying to find your way out of a forest. A sign pointing right and a distant sound from the left provide evidence for different directions. If choosing right leads to success, the sign would be updated more strongly because it supported the chosen action. Yet if the sound is reactivated when the outcome is received, its representation may also be updated more strongly than expected from its contribution to the choice alone. More generally, even for the same choice and outcome, different cues can contribute unequally to the decision and be differentially represented during feedback, causing their predictive values to be updated to different degrees. For these reasons, the brain may not treat credit assignment and learning-rate control separately, but instead couples what is learned with how strongly it is updated.

Rather than merely a byproduct of biological constraints on learning and plasticity, we hypothesize this coupling might serve a functional role by making learning more efficient and enhancing discrimination between competing predictive cues. When multiple predictive cues are present, less informative cues can contribute more strongly to prediction errors and therefore undergo larger value changes, progressively reducing differences in their learned predictive values. We postulated learning rates are structured by cue-choice-outcome interactions arising from the coactivation of task-relevant representations during naturalistic learning, thereby linking credit assignment and adaptive learning-rate control.

To test these, we asked human participants to perform a multi-cue probabilistic learning task in which they predicted one of two weather outcomes (rainy vs. windy) based on four visual cues and received reward feedback after each choice. Simultaneously, we also tracked eye gaze throughout each trial to infer how attention, guided by the value of each cue (i.e., value-guided attention), was allocated during decision making and learning. This design allowed us to dissociate choices from the cues that predicted them, identify which representations were preferentially processed, and determine how cues, choices, and reward outcomes interact to determine learning. Critically, we also manipulated these processes by changing cue saliency at the time of feedback to test its causal influence on learning.

By constructing and fitting a set of computational models to participants’ choices, we found that learning rates were systematically influenced by the alignment between cues, choices, and reward outcomes. Specifically, on rewarded trials, learning was biased toward cues that support the chosen option, whereas on unrewarded trials, learning was biased toward cues that oppose it. Consistent with our hypothesis, this cue-choice congruency-dependent updating improved discrimination between competing cues, robustness to noise, and overall performance. Moreover, trial-by-trial gaze patterns mirrored the learning-rate asymmetry, and individual differences in dwell time on different cues predicted corresponding differences in learning rates. Finally, manipulating cue saliency altered both gaze patterns and learning-rate asymmetries as predicted. Together, our findings provide a unified account of previously reported learning asymmetries and suggest that attention-guided coactivation of task-relevant representations links credit assignment and learning rates to enhance cue discriminability and learning robustness.

## Results

### Participants successfully learned individual cue-outcome associations

To investigate how learning rates depend on cue-choice-outcome interactions, participants performed a multi-cue probabilistic learning task in which they predicted one of two weather outcomes (windy vs. rainy) based on four visual cues (simple shapes, with possible repetitions). After making their choice, they received reward feedback, allowing them to learn the associations between individual cues and choice outcomes (**Fig. 1a–c**). In addition, the task included blocks of estimation trials in which participants used the keyboard to report their estimates of the probability that a given cue or combination of cues predicted a windy or rainy outcome, with single-cue estimates obtained in the first three blocks and multi-cue (two-and four-cue) estimates collected in the final block.

**Figure 1.**
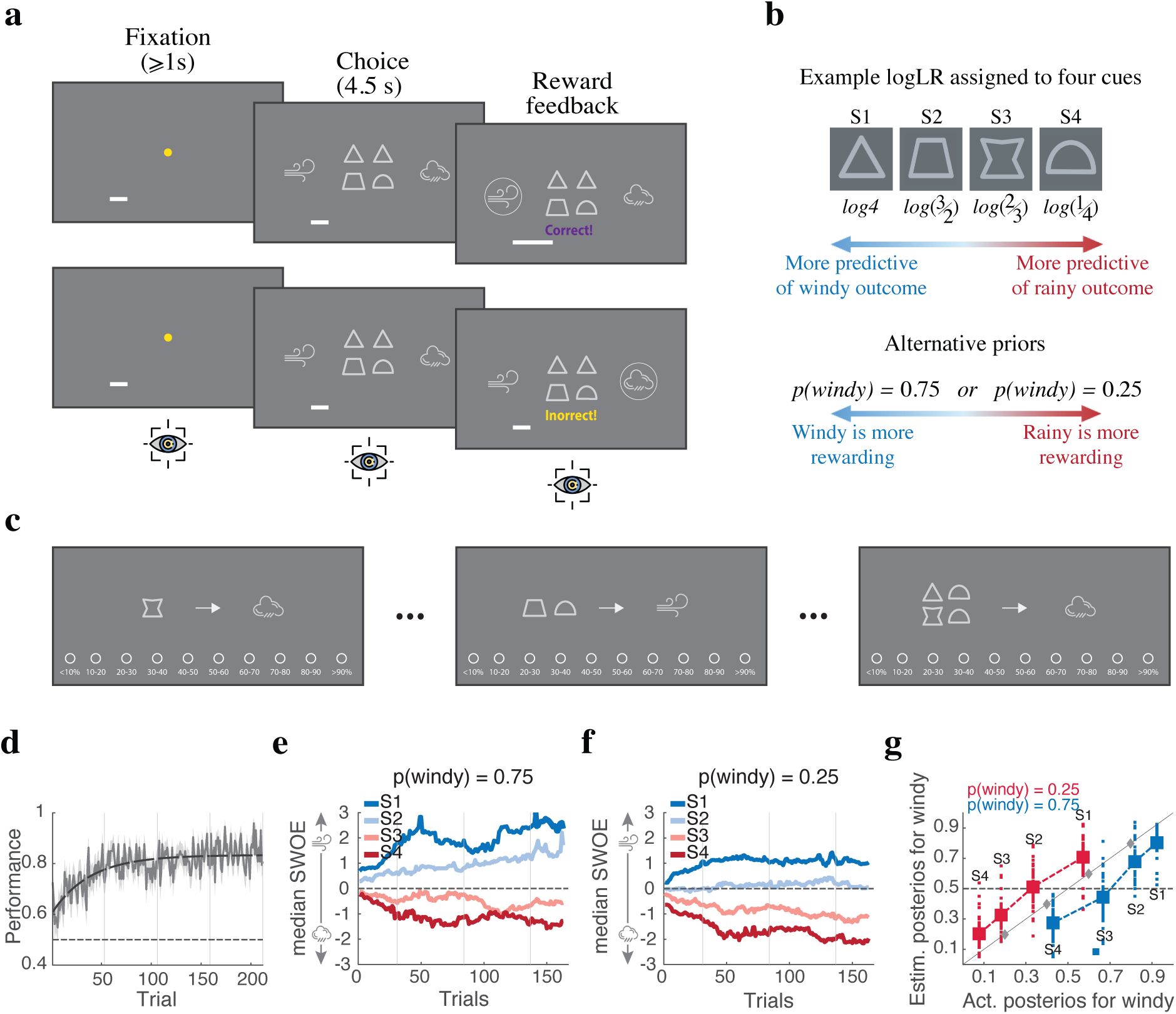
Experimental paradigm and evidence for successful learning of cue-outcome associations. (**a**) Timeline of a choice trial. Each trial began with fixation on a central yellow circle, followed by presentation of four visual cues and two choice options (‘windy’ vs. ‘rainy’). The selected option was indicated with a circle, after which reward feedback (‘correct’ or ‘incorrect’) and an updated reward bar were shown. Participants’ gaze was recorded continuously throughout the experiment. (**b**) Example cues and their evidence (logLR) for the windy option. Participants performed the task with p(windy)=0.25 or 0.75. (**c**) Schematic of example estimation trials. On each trial, participants used the keyboard to select from 10 values to report the probability that a given cue or combination of cues predicted a windy or rainy outcome. In the first three blocks, estimates were based on single cues; in the final block, these were followed by estimates for combinations of two cues and then four cues. (**d**) Time course of participants’ performance, defined as the proportion of trials in which the correct option (option with higher posterior probability) was chosen. The plot shows average performance across participants during choice trials. The shaded area represents the SEM, and vertical bars indicate the positions of estimation blocks. The black curve shows an exponential fit to performance. (**e,f**) Time courses of the median subjective weights of evidence (SWOEs) provided by the four cues, shown separately for participants with prior probabilities of 0.75 (e) and 0.25 (f). Overall, SWOEs were biased toward the higher-prior option. (**g**) Estimated posterior probability of the windy option for individual cues as a function of the true posterior probability, shown separately for participants with priors of 0.25 and 0.75. Each dot represents the average estimate from an individual across four estimation blocks, while solid squares indicate the group average across all participants. Gray diamonds denote the evidence associated with each cue, and dashed lines are shown for visual guidance. Overall, participants overestimated posteriors for cues favoring the lower-prior option.

Critically, individual cues differed in their predictive power for the rainy and windy outcomes, while one outcome had a higher prior probability for each participant (the higher-prior option), making it more often the correct choice (see **Methods**). This allowed us to examine how participants learned the evidence provided by each cue, as well as differences in reward expectations associated with the two choice options, which we refer to as the higher-prior and lower-prior options. This design ultimately enabled us to reveal how prediction, expectation, choice, and reward outcomes jointly influence learning. We also tracked participants’ eye movements throughout each trial to determine how attention to different cues and choice options evolved over time, allowing us to assess how attentional allocation influenced decision making during deliberation and value updating during reward feedback.

Overall performance was significantly above chance as participants selected the correct option (i.e., the one with higher posterior probability) on most trials (mean = 0.79, t(72) = 32.29, p = 1.41 × 10^−44^, d = 3.78). Moreover, the time course of performance indicated that participants learned the task rapidly, with performance approaching asymptote after ∼70 trials (**Fig. 1d**). To capture the evolution of learned cue– outcome associations, we used logistic regression with a sliding window of 45 trials centered on each trial to estimate the subjective weight of evidence (SWOE) each cue provided for the windy (vs. rainy) option. We found that SWOEs rapidly diverged, indicating that participants learned the predictive values of individual cues (**Fig. 1e,f**). Moreover, although the relative ordering of cue values remained consistent with their objective evidence, SWOEs were biased toward the higher-prior option, being shifted toward more positive values when p(windy)=0.75 and toward more negative values when p(windy)=0.25.

Analysis of estimation trials showed that the ordering of estimated posteriors for individual cues matched the evidence provided by those cues (**Fig. 1g**). However, estimates were systematically biased toward the lower-prior option. When p(windy)=0.25, estimated posteriors for the windy option exceeded the true values (S1: t(35)=6.07, p=6.23×10^−7^, d=1.01; S2: t(35)=7.71, p=4.69×10^−9^, d=1.28; S3: t(35)=7.09, p=2.85×10^−8^, d=1.18; S4: t(35)=6.40, p=2.27×10^−7^, d=1.06), whereas when p(windy)=0.75, they fell below the true values (S1: t(36)=-7.49, p=7.45×10^−9^, d=-1.23; S2: t(36)=-7.09, p=2.45×10^−8^, d=-1.16; S3: t(36)=-8.59, p=3.01×10^−10^, d=-1.41; S4: t(36)=-8.49, p=3.99×10^−10^, d=-1.39). Consistent with previous findings (Soltani et al., 2016), this bias can be explained by a model in which posterior beliefs are represented by cue-specific estimates for the two choice options. Because multiple cues jointly contribute to behavior, each cue carries only part of the decision signal, yielding cue-specific posterior estimates that are smaller than the full Bayesian posterior integrating prior and cue evidence.

### Learning rates were shaped by cue-choice alignment and reward outcome

To test our hypothesis that learning rates are jointly structured by cue-choice alignment and reward outcome, we built multiple computational models to fit choice behavior on a trial-by-trial basis. This included a single “baseline” model that assumed updating was symmetrical across reward outcomes and across cues, so that a single learning rate governed all updates. All remaining models incorporated some form of learning-rate asymmetry. The simplest allowed separate learning rates for rewarded and unrewarded outcomes (reward/no reward model), independent of the cues used to make decisions. We also included two cue-dependent models. The first allowed learning rates to vary across cues depending on whether they supported the higher-prior or lower-prior option, with separate learning rates for rewarded and unrewarded trials (cue-prior model). This model captures learning structured by beliefs about globally more rewarding options. The second model instead tied learning rates to cue-choice alignment/congruency on each trial (cue-choice congruency model), allowing rewarded and unrewarded outcomes to differentially update cues that support versus oppose the chosen option. Thus, whereas the first cue-dependent model tested whether learning is structured by beliefs about the globally more rewarding option, the second tested our hypothesis that learning rates are jointly shaped by cue-choice congruency and reward outcome. Finally, to test whether mechanisms beyond learning-rate asymmetry contribute to choice behavior, we extended all non-baseline models with variants that also incorporated asymmetric weighting during decision making. These asymmetries could occur either between choice options (higher-vs. lower-prior; option weighting) or between predictive cues (those favoring the higher-vs. lower-prior option; cue weighting). These variant allowed us to dissociate effects on learning from differential evidence weighting during choice.

Using Bayesian model selection (BMS) (Rigoux et al., 2014; Stephan et al., 2009), we found that the cue-choice congruency model provided the best fit both at the individual and group levels (posterior probability = 0.37; PXP = 0.58; **Fig. 2a**). The second-best model additionally incorporated differential weighting of cues during decision making, alongside cue-choice congruency-dependent learning (posterior probability = 0.35; PXP ≈ 0.41; **Fig. 2a**). These results underscore that explaining participants’ choice behavior requires learning to depend jointly on cue-choice congruency and reward outcome. We validated these results through model recovery, showing that the best-fitting model could be reliably distinguished from alternative models (**Fig. S1a–d**).

**Figure 2.**
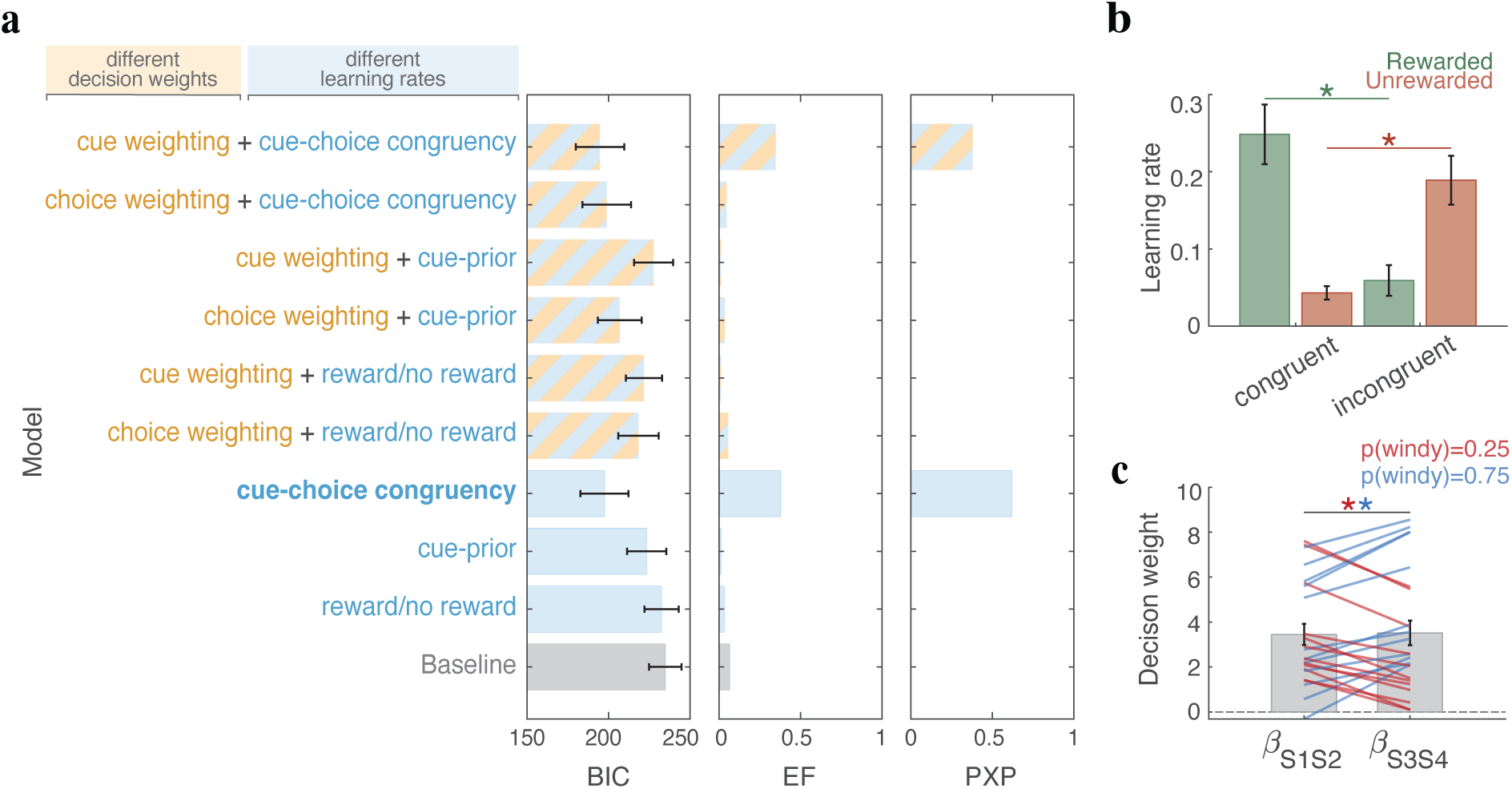
Choice behavior was best fit by a model incorporating cue-choice congruency-dependent learning. (a) Model comparison using Bayesian Information Criterion (BIC), estimated frequency (EF), and protected exceedance probability (PXP). Lower BIC values indicate better fit, whereas higher EF and PXP indicate stronger evidence for a model. The cue-choice congruency model (bold), that allows rewarded and unrewarded outcomes to differentially update associations for cues that support versus oppose the chosen option, provides the best overall fit. (b) Estimated learning rates from the best-fitting model for cues congruent and incongruent with the chosen option, shown separately for rewarded and unrewarded trials. Error bars represent the SEM across participants. Asterisks denote significant differences (Wilcoxon signed-rank test, p < 0.05). (**c**) Estimated decision weights for two sets of cues supporting the higher-prior and lower-prior options, shown for participants best fit by the second-best model (cue-choice congruency with differential cue weighting). Red and blue asterisks indicate significant differences between the two weights (Wilcoxon signed-rank test, p < 0.05) for p(windy)=0.25 and p(windy)=0.75, respectively. Each line represents an individual participant, and error bars denote the SEM.

Analysis of learning rates from the best-fitting model further revealed a systematic dependence on the cue-choice-outcome interactions. On rewarded trials, learning rates were higher for cues supporting the chosen option (choice-congruent cues; S1 and S2 for windy, S3 and S4 for rainy) than for cues opposing it (choice-incongruent cues; S1 and S2 for rainy, S3 and S4 for windy) (Wilcoxon signed-rank test: z = 5.14, p = 2.62 × 10^−7^, d = 0.60; **Fig. 2b**). In contrast, on unrewarded trials, learning rates were higher for cues opposing the chosen option than for those supporting it (z = 4.24, p = 2.16 × 10^−5^, d = 0.49; **Fig. 2b**). Importantly, this pattern was independent of which outcome was more frequently rewarded (**Fig. S2a**) and was observed when fitting participants according to their individually best-fitting models (**Fig. S2b,c**). Parameter recovery analysis showed that estimated parameters closely matched the true values, with strong correlations and minimal systematic bias (**Fig. S1e–f**). These results demonstrate that learning asymmetry extends to the level of predictive cues: rewarded outcomes preferentially enhance learning from cues supporting the chosen option, whereas unrewarded outcomes enhance learning from cues opposing it.

Analysis of the estimated decision weights in the cue-choice congruency model with differential cue weighting revealed that participants best described by this model placed greater weight on cues predictive of the lower-prior option than on cues predictive of the higher-prior option (Wilcoxon signed-rank test; p(windy)=0.25: z = 3.05, p = 4.88 × 10^−4^, d=0.88; p(windy)=0.75: z = 3.05, p = 4.88 × 10^−4^, d=0.88; **Fig. 2c**). This pattern may reflect differences in how often cues are encountered. For a given prior, cues supporting the higher-prior option appear more often than cues predicting the alternative. The stronger weighting of cues supporting the lower-prior option may therefore reflect greater sensitivity to less frequently encountered evidence.

As one of several control analyses, we tested a model in which participants infer the opposite outcome for the unchosen option (i.e., reward for one option implies no reward for the other, and vice versa) and update it with a separate learning rate. However, the simpler model with differential learning rates for rewarded and unrewarded outcomes (reward/no reward model) provided a better fit, indicating that asymmetric updating of the unchosen option did not improve fit of choice data and explanatory power (**Fig. S3**). In addition, we also compared the best-fitting (cue-choice congruency) model to an alternative in which learning rates depended on the congruency between each cue and participants’ predicted choice (cue-prediction congruency model), rather than their actual choice on each trial. To that end, we used the sum of the difference in synaptic strengths for the presented cues, as a measure of the option predicted by those cues, instead of the participant’s actual choice to determine congruency. Model selection favored the cue-choice congruency model over this cue-prediction model, indicating that the observed learning asymmetry reflects alignment between cues and actual choices, rather than predictions or beliefs (**Fig. S4**).

### Gaze patterns predicted cue-choice congruency-dependent learning

Model fits to choice behavior revealed reward-dependent learning asymmetries that varied with cue-choice congruency; that is, whether a cue provides evidence supporting or opposing the chosen option. These models build on reward-dependent Hebbian plasticity (Soltani et al., 2016; Soltani & Wang, 2006, 2010), in which learning occurs when representations of a cue and an outcome are active at the same time, while reward signals such as dopamine determine whether the association between them is strengthened or weakened. Such coactivation may accumulate during deliberation via an eligibility trace (Frémaux & Gerstner, 2016; Izhikevich, 2007; Shindou et al., 2019; Shouval & Kirkwood, 2025) or arise at the time of reward feedback. Because fixation enhances neural responses to attended stimuli (Desimone & Duncan, 1995; J. H. Reynolds & Chelazzi, 2004), and gaze is systematically influenced by ongoing choice processes (Anderson, 2013; Cavanagh et al., 2014; Krajbich et al., 2010), eye movements can provide an indirect measure of which cue and choice representations are preferentially processed during different phases of a trial.

To test this, we quantified the proportion of time participants spent on each cue and choice option relative to all task elements within each epoch, averaged across trials (**Methods**). General analyses of fixations on cues and choice options indicate that gaze systematically reflects decision-making processing and learned cue-outcome associations (**Fig. S5; Supplementary Note 1**). More specifically, during the choice epoch, participants preferentially directed their gaze toward the higher-prior option (**Fig. S5b**) and toward cues that supported their eventual choice on a given trial (**Fig. S5c**), but showed no systematic preference for one cue group over another (**Fig. S5a**). During the feedback epoch, participants continued to preferentially fixate the higher-prior option (**Fig. S5e**), albeit less strongly than during the choice epoch, and showed a modest tendency to fixate cues supporting it (**Fig. S5d**). They also spent significantly more time fixating cues that supported the chosen option than cues that opposed it (**Fig. S5f**). Together, these findings suggest that gaze reflects both learned evidence and choice-related information across time. We therefore next examined whether gaze behavior mirrored the cue–choice congruency effect on learning by quantifying dwell time as a function of cue–choice congruency.

We found that during the feedback epoch of rewarded trials, participants preferentially fixated cues congruent with the chosen option compared to those that were incongruent (Wilcoxon signed-rank test; z = 2.62, p = 0.008, d = 0.30; **Fig. 3a,b**). This pattern reversed on unrewarded trials, with participants dwelling more on cues incongruent with the chosen option (z = 2.12, p = 0.033, d = 0.25; **Fig. 3a,b**). Importantly, a linear mixed-effects regression revealed a significant interaction between reward feedback and cue-choice congruency (b = 0.161, SE = 0.031, t(288) = 5.09, p = 6.43 × 10^−7^), indicating that the effect of congruency on gaze depended on whether the choice was rewarded. There were also significant main effects of feedback (b = −0.093, SE = 0.022, t(288) = −4.14, p = 4.39 × 10^−5^) and congruency (b = −0.071, SE = 0.027, t(288) = −2.61, p = 0.009). Together, these results show that during the feedback epoch, participants preferentially attended to congruent cues following reward and to incongruent cues following non-reward, mirroring cue-choice congruency-dependent learning rates.

**Figure 3.**
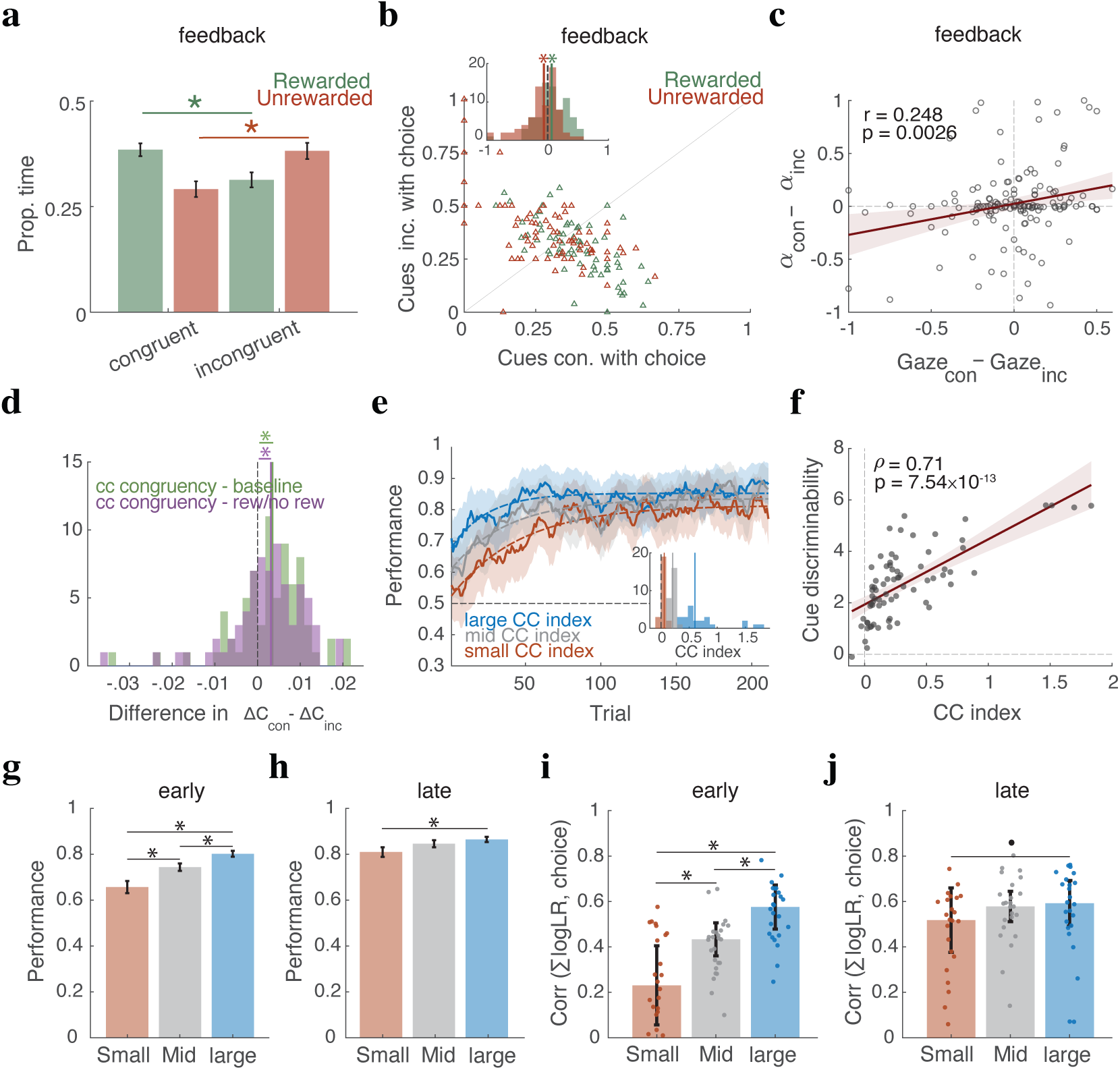
Gaze predicts congruency-dependent learning that enhances robustness and performance. (**a–c**) Differential gaze to congruent vs. incongruent cues during the feedback epoch and its link to learning-rate asymmetry. Plots show (a) mean proportion of dwell time on each cue type for rewarded and unrewarded trials (mean ± SEM; asterisk shows Wilcoxon signed-rank test, p < 0.05), (b) scatter plots and distributions of the difference (congruent − incongruent) for each reward feedback, and (c) across-participant correlation between differential processing and learning-rate asymmetry. Medians and asterisks indicate significant deviations from zero, and the regression line is shown in red, with the correlation coefficient and p-value reported. (**d**) Difference in trial-by-trial cue-specific updates between cues congruent and incongruent with the chosen option, comparing the cue-choice congruency model with two control models (baseline and reward/no reward models). Dashed and solid vertical lines indicate zero and medians, respectively. (**e**) Time course of performance, shown separately for participants grouped by their cue-choice congruency (CC) index: low (red), middle (gray), and high (blue). Mean performance is smoothed with a 10-trial moving average, with shaded regions indicating SEM. Dashed curves represent exponential fits to the group-level performance. (**f**) Cue discriminability, measured as (S1′−S4′) + (S2′−S3′) from participants’ SWOEs, plotted as a function of each participant’s CC index. Higher values indicate greater separation between cues providing opposing evidence. The red line and shaded band show the linear regression fit and its 95% confidence interval. (**g,h**) Mean performance during early (trials 1–70) and late (final 70 trials) phases of the experiment, shown separately for participants grouped by CC index. Error bars represent ± SEM. During the early phase, participants with higher CC indices showed significantly better performance, whereas group differences were reduced in the late phase. **(i,j)** Across-participant correlation between the summed objective weights of evidence (sum logLR) carried by the presented cues and choice during early (trials 1–70) and late (final 70 trials) phases of the experiment, shown separately for participants grouped by CC index. Bars show medians, error bars indicate half the interquartile range, and dots represent individual participants; filled circles (•) denote marginal significance (Wilcoxon test, p = 0.05). Higher values indicate stronger alignment between objective cue evidence and choice.

Considering dwell time as a proxy for cue-and choice-related processing and activation of task-relevant representations, we next asked whether it is linked to the observed asymmetry in learning rates. To this end, we quantified learning-rate asymmetry as the difference between congruent and incongruent cues (αcon-α_inc_), separately for rewarded and unrewarded trials. Across participants, asymmetry in learning rates was significantly correlated with differences in dwell time on congruent versus incongruent cues during the feedback epoch (Pearson correlation; r = 0.25, p = 0.002; **Fig. 3c**). This demonstrates that the cue-choice congruency effect on learning is closely linked to how cue and choice-outcome information are processed during reward feedback.

We also tested whether cue-choice-outcome interactions influenced subsequent decision making by biasing cue processing on the next trial, but found no evidence for such effects (**Fig. S6a–c**). Similarly, we found no differences in dwell time between congruent and incongruent cues during the choice epoch, nor any relationship between dwell time and the learning asymmetry (**Fig. S6d–f**). These suggest that differential learning is mainly influenced by the pattern of dwell time after the presentation of reward feedback, and not during choice, highlighting the modulation of learning by value-guided overt attention.

Finally, among participants exhibiting cue-weighting effects, differential dwell time on congruent versus incongruent cues during the choice epoch did not predict the magnitude of cue weighting (**Fig. S7a,b**). Consistent with this, a mixed-effects logistic regression revealed no evidence that dwell time on cues improved choice prediction beyond a null model (**Table S1**). Nonetheless, dwell time on choice options was informative: trial-by-trial differences in dwell time on the two options significantly improved the prediction accuracy relative to a null model (**Table S1**). Thus, although attention toward the choice options explained some variance in participants’ choices, attention toward cues did not predict variability in the weights of cue groups on choice.

Together, these results suggest that value-guided overt attention primarily influences learning rather than decision making by differentially activating cue representations during feedback: choice-congruent cues on rewarded trials and choice-incongruent cues on unrewarded trials. This activation pattern, in turn, influences learning-rate asymmetries by increasing learning rates for congruent cues on rewarded trials and for incongruent cues on unrewarded trials.

### Cue-choice congruency effect does not require unequal priors

We also tested whether the observed cue-choice congruency learning asymmetry depends on unequal priors, which could introduce asymmetries between choice options and their associations with cues. To this end, we conducted a control experiment with equal probabilities for rainy and windy outcomes in a separate group of participants (N = 32). Model comparison showed that the cue-choice congruency model again provided the best overall fit to the choice data (**Fig. S8a**). Consistent with our main findings, estimated learning rates exhibited the same pattern: higher learning rates for congruent than incongruent cues on rewarded trials, and higher learning rates for incongruent than congruent cues on unrewarded trials (**Fig. S8b**). These results demonstrate that the cue-choice congruency effect extends beyond conditions with unequal priors and support our hypothesis that learning-rate asymmetries are influenced by neural activation driven by value-guided overt attention.

### Cue-choice congruency effect increased robustness to noise and improved performance

Given the presence of cue-choice congruency-dependent learning and its link to value-guided attention, a key question is what functional role this learning serves. We hypothesized that learning-rate asymmetries preserve discriminability among cues’ learned predictive values, which would otherwise diminish following repeated updates associated with the more common outcome.

The rationale is that, when there are competing predictive cues, cues that more strongly support the chosen option contribute more to the predicted outcome and therefore generate smaller prediction errors, whereas cues opposing the chosen option generate larger prediction errors (or contribute more strongly to the overall prediction error). Consequently, when all presented cues are updated at the same rate, opposing cues undergo larger value changes than supporting cues, progressively reducing the separation between learned cue predictive values over time. We therefore predicted that asymmetric learning rates, favoring cues congruent with the chosen option following the more common outcome (reward) and incongruent cues following the less common outcome (no reward), would counteract this effect and maintain greater discriminability among predictive cues.

To test this hypothesis, we examined whether individual differences in learning-rate asymmetry predicted cue discriminability, quantified as (S1′−S4′) + (S2′−S3′) based on participants’ SWOEs (S1′–S4′ denote the cues reordered from most to least predictive of the higher-prior option). We quantified learning-rate asymmetry using a cue-choice congruency (CC) index, defined as the average difference between learning rates for congruent and incongruent cues on rewarded trials and for incongruent and congruent cues on unrewarded trials, normalized by the inverse-temperature parameter (see **Methods**). We found that participants with higher CC indices exhibited greater discriminability between cues providing opposing evidence (Spearman’s ρ = 0.71, p = 7.54 × 10^−13^; **Fig. 3f**), consistent with the idea that cue-choice congruency-dependent learning enhances the separation of predictive cue values.

To assess how learning-rate asymmetry shaped trial-by-trial updating, we used cue-specific estimates for the two choice options (represented by separate synaptic strengths; see Eqs. 5–7) to quantify how much the predictive evidence associated with each cue changed after each trial. We compared these updates across the best-fitting model and two control models: the baseline model with a single learning rate and the model with separate learning rates for rewarded and unrewarded outcomes. If our hypothesis is correct, the best-fitting model should produce larger updates than the control models for cues congruent with the chosen option after reward and for cues incongruent with the chosen option after no reward.

Consistent with our hypothesis, the cue-choice congruency model produced larger trial-by-trial differences in updating between cues that supported versus opposed the chosen option than either control model (Wilcoxon signed-rank test: cue-choice congruency vs. single-rate model, z = 3.66, p = 2.48 × 10^−4^, d = 0.42; cue-choice congruency vs. reward/no-reward model, z = 3.05, p = 0.002, d = 0.35; **Fig. 3d**). These results indicate that congruency-dependent learning-rate asymmetry helps preserve differences in the predictive values of competing cues over learning. Whether this preservation also improves learning efficiency and behavioral performance, however, remains an open question.

To answer these, we compared performance across participants with different levels of cue-choice congruency. Dividing participants into tertiles based on their CC index revealed that the high-CC group achieved both higher asymptotic performance (0.853) and faster learning (τ = 23.4) than the mid-CC (0.836, τ = 38.0) and low-CC groups (0.814, τ = 46.2) (**Fig. 3e**). Comparing early and late phases of the experiment, we found that during the early phase, participants with higher CC indices showed significantly better performance than those with lower CC indices (Wilcoxon rank-sum tests: large vs. small, z = 3.96, p = 7.31 × 10^−5^, d = 0.57; large vs. mid, z = 2.54, p = 0.0109, d = 0.36; mid vs. small, z = 2.42, p = 0.0153, d = 0.34; **Fig. 3g**). In the late phase, the advantage of the high-CC group over the low-CC group remained significant (z = 1.96, p = 0.049, d = 0.28), whereas differences between the other groups were no longer significant (large vs. mid: z = 0.59, p = 0.553, d = 0.08; mid vs. small: z = 1.14, p= 0.253, d = 0.16; **Fig. 3h**). Critically, comparing the cue-choice congruency effect between the two halves showed that the effect was stronger in the second half of the experiment (**Fig. S2d,e**), indicating that cue-choice congruency-dependent updating strengthens over the course of learning. Therefore, more similar performance later in the experiment was mainly due to ceiling effect. Together, these results suggest that stronger cue-choice congruency is associated with both faster and more robust learning.

Finally, if the performance differences reflect better discrimination of cue predictive values, they should also be accompanied by a stronger relationship between choice and objective cue evidence. To test this, we computed the correlation between choice and the summed objective weights of evidence (sum logLR) of the presented cues during early (trials 1–70) and late (final 70 trials) phases. In the early phase, groups differed significantly, with higher CC indices associated with stronger alignment between evidence and choice (Wilcoxon rank-sum tests: large vs. small, z = 4.52, p = 6.01 × 10^−6^, d = 0.65; large vs. middle, z = 3.53, p = 4.15 × 10^−4^, d = 0.50; middle vs. small, z = 2.47, p = 0.0135, d = 0.35; **Fig. 3i**). In the late phase, this pattern was weaker perhaps due to ceiling effect: only the difference between high-and low-CC groups remained marginally significant (z = 1.88, p = 0.059, d = 0.27; **Fig. 3j**), with no differences between other groups. These findings were corroborated by GLM analyses showing that cue-choice congruency asymmetry predicted both performance (**Table S2**) and sensitivity to objective evidence (**Table S3**). Together, these results suggest that stronger cue-choice congruency asymmetry enhances performance by increasing sensitivity to objective evidence, particularly early in learning, with a reduced but still detectable effect later on.

### Causal manipulation of cue fixation and processing alters learning

Our results thus far suggest that value-guided overt attention during feedback drives cue-choice congruency-dependent learning and enhances discrimination among predictive cues. They also provide an explanation for the observed pattern of learning-rate asymmetry. Specifically, higher learning rates for congruent than incongruent cues on rewarded trials are expected because neural populations selective for congruent cues are more strongly activated during both choice and feedback, due to their contribution to decision making and their preferential activation by value-guided attention during feedback. More puzzling is the higher learning rates for incongruent than congruent cues on unrewarded trials. If reward omission simply reversed plasticity from potentiation to depression through reduced dopamine release, stronger depression would be expected for congruent cues instead. One possible explanation is that increased fixation on incongruent cues during feedback enhances activation of their neural representations, leading to stronger synaptic depression (larger α^−^). We next tested this possibility by causally altering cue fixation during feedback.

To that end, a subset of participants (N = 37) completed an additional session in which the luminance of one predefined low-informative cue (S2 or S3) was transiently increased during reward feedback (**Fig. S9a,b**), allowing within-subject comparisons with and without the saliency manipulation. For some participants, the manipulated cue was more predictive of the higher-prior option, whereas for others it was more predictive of the lower-prior option, making the salient cue more or less likely to align with their choices, respectively. Consequently, participants were divided into two groups using a saliency-congruency index (see **Methods**). Participants whose salient cue was predominantly choice-congruent were assigned to the congruent-saliency manipulation group, whereas those whose salient cue was predominantly choice-incongruent formed the incongruent-saliency manipulation group.

We predicted that learning asymmetry would be disrupted in the congruent-manipulation group because luminance increase and subsequent fixation (exogenous overt attention) would more frequently activate congruent than incongruent cues during feedback, thereby reducing the differential activation of incongruent cues normally observed on unrewarded trials. In contrast, learning asymmetry was expected to persist in the incongruent-manipulation group because this manipulation would less frequently disrupt the stronger activation of incongruent cues during feedback, while congruent cues still benefit from enhanced activation during decision making.

We found that the saliency manipulation successfully altered overt attention during the feedback epoch. In both groups, participants spent significantly more time fixating the salient cue than the non-salient cues (Wilcoxon signed-rank test; congruent manipulation: z = 3.24, p = 0.001, d = 0.72; incongruent manipulation: z = 3.05, p = 0.002, d = 0.74; **Fig. 4a**). Moreover, under congruent manipulation, participants fixated S2′ more than S3′ (Wilcoxon signed-rank test: z = 3.21, p = 0.001, d = 0.71; **Fig. 4b**), whereas under incongruent manipulation, they fixated S3′ more than S2′ (z = 2.76, p = 0.005, d = 0.67; **Fig. 4b**). This manipulation disrupted the cue-choice congruency pattern in gaze, as revealed by fixation times analyzed as a function of cue-choice congruency (compare **Fig. 4c,d** with **Fig. S5f**).

**Figure 4.**
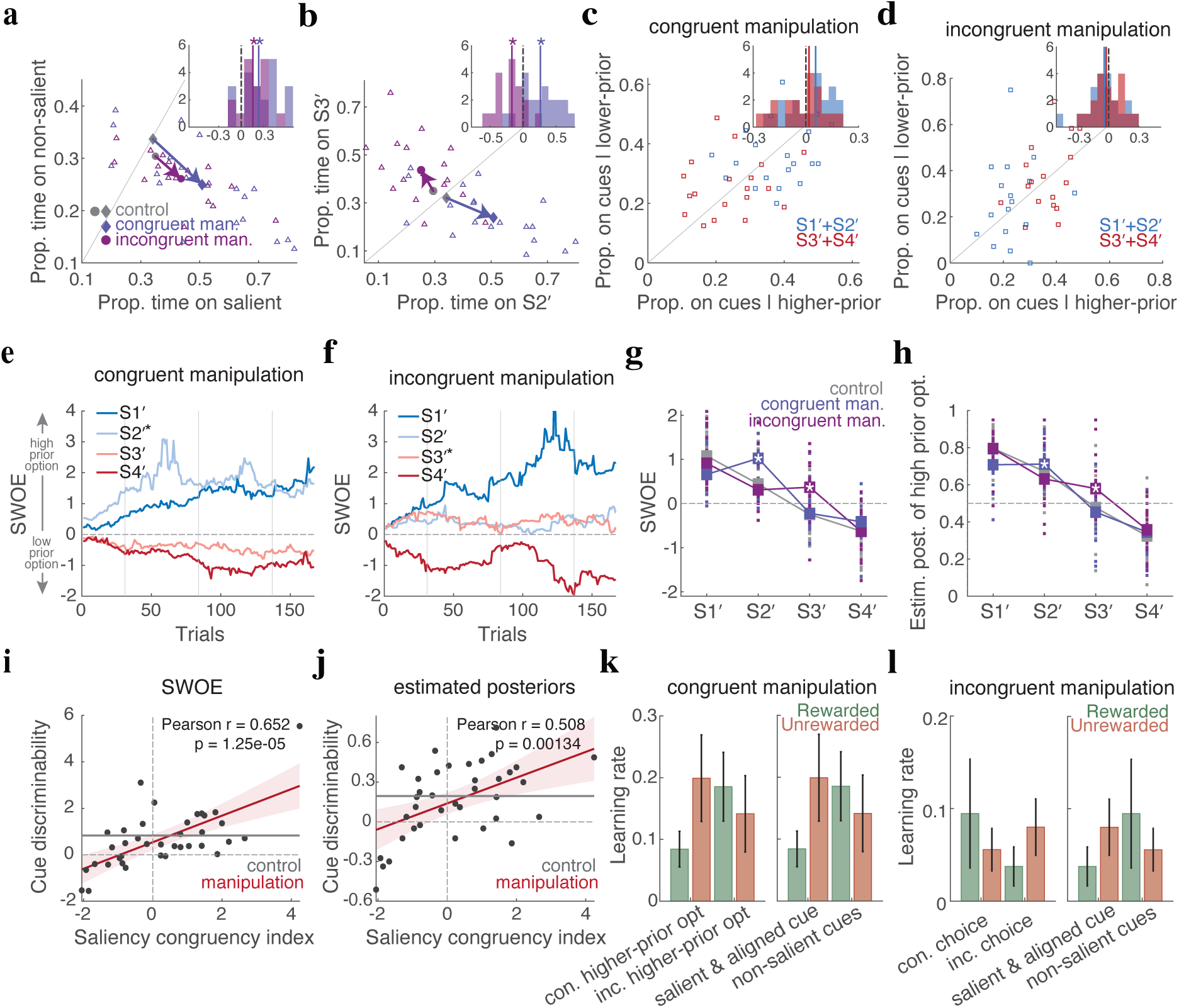
Saliency manipulation during feedback alters learning and cue discriminability. (**a,b**) Changes in gaze due to saliency manipulation during the feedback, shown separately for participants whose salient cue was more often choice-congruent or choice-incongruent: (a) gaze time on the salient cue versus the mean of non-salient cues; (b) gaze time on S2′ versus S3′. S2′ and S3′ denote cues more predictive of the higher-and lower-prior options, respectively. Larger markers denote group centroids (see legend), and arrows connect the control centroid to each manipulation-group centroid. Histograms show within-subject differences between the y-and x-axis variables for each group; solid vertical lines indicate median differences, and stars denote significant deviations from zero (p < 0.05). In both groups, participants spent more time viewing the salient cue than the non-salient cues (a). Saliency manipulation also biased dwell time between S2′ and S3′: the congruent-manipulation group fixated on S2′ more than S3′, whereas the incongruent-manipulation group showed the opposite pattern (b). (**c,d**) Mean proportion of dwell time on cues supporting the chosen vs. unchosen option during the feedback epoch, separately for each group. Histograms show differences, with median differences indicated, and stars denote significant deviations from zero. (**e,f**) Time courses of median SWOEs for the four cues, shown separately for each group. Positive (negative) SWOE values indicate weighting toward the higher-prior (lower-prior) option. The salient cue in each group is indicated by a star in the legend. (**g,h**) Median SWOEs (g) and posterior estimates from the estimation sessions (h) for individual cues, shown for the congruent-manipulation group, the incongruent-manipulation group, and the same participants in the control condition without saliency manipulation. The salient cue in each manipulation group is marked with a star, and small squares indicate individual participants’ values. (**i,j**) Discriminability between the two least informative cues (S2′ and S3′) plotted against each participant’s saliency-congruency index, measured using SWOEs (i) and posterior estimates (j). The solid red line shows the linear regression fit, with the shaded band indicating the 95% confidence interval. The solid gray line indicates baseline cue discriminability for the same participants in the absence of manipulation. (**k,l**) Estimated learning rates of the winning model for participants best fit by the cue-prior model with differential option-weighting in the congruent-manipulation group (k) and by the cue-choice congruency model in the incongruent-manipulation group (l). Left panels show the original learning rates; right panels reorganize them according to whether the cue group more often contained the salient or non-salient cues. Error bars indicate SEM.

Moreover, we found that saliency manipulation affected behavior differently depending on the congruency of the manipulated cue with choice on a given trial. In the congruent-manipulation group, where the salient cue more often supported the chosen option, performance did not differ from control during the early phase (Wilcoxon rank-sum test: z = −1.27, p = 0.203, d = −0.28) but was significantly worse during the late phase (z = −2.48, p = 0.012, d = −0.55; **Fig. S9c,e,f**). Alignment between objective evidence and choice was also reduced in both phases (**Fig. S9g,h**). In contrast, the incongruent-manipulation group, in which the salient cue more often supported the unchosen option, showed no evidence of impaired performance or reduced evidence-choice alignment relative to control in either phase (Performance: early: z = −0.75, p = 0.452, d = −0.18; late: z = −1.11, p = 0.264, d = −0.27; **Fig. S9d–f**; evidence-choice alignment: early: z = −1.77, p = 0.075, d = −0.43; late: z = −1.49, p = 0.135, d = −0.36; **Fig. S9g,h**).

Analysis of SWOE time courses revealed that, in both manipulation groups, the salient cue rapidly became more strongly weighted as evidence for the higher-prior option. In the congruent group, the SWOE of S2′ exceeded that of S1′, indicating that participants treated S2′ as more predictive of the higher-prior option than the objectively strongest cue (**Fig. 4e,g**). In the incongruent group, the SWOE of S3′ shifted from negative to positive, indicating that participants treated a cue predictive of the lower- prior option as evidence for the higher-prior option (**Fig. 4f,g**). The same pattern was reflected in participants’ reported posterior estimates (**Fig. 4h**), even though saliency manipulation occurred only during choice trials, suggesting that the manipulation altered learning itself. These effects could arise either directly from the luminance increase enhancing activation of the manipulated cue, or indirectly through the increased overt attention elicited by the luminance manipulation.

To test the impact of saliency manipulation on cue discriminability, we examined whether the saliency-congruency index used to divide participants into the two groups predicted discriminability between S2′ and S3′. The saliency-congruency index strongly predicted discriminability computed from both SWOEs (r = 0.65, p = 1.25 × 10^−5^; **Fig. 4i**) and participant estimates (r = 0.51, p = 0.001; **Fig. 4j**). Thus, the more often saliency aligned with choice, the more strongly participants discriminated the two cues in the correct direction, whereas predominantly incongruent saliency reversed this pattern. This demonstrates that increasing cue luminance can enhance its predictive value and/or its influence on decision making.

To tease apart these two possibilities, we next fit participants’ choices using our computational models and found that learning-rate asymmetry was primarily disrupted in the congruent-saliency group. For this group, the best-fitting model was the cue-prior model with differential choice weighting, in which learning rates depended on whether cues supported the higher-or lower-prior option (S1 and S2 vs. S3 and S4) rather than on the participant’s choice (**Fig. S10**). Moreover, the estimated learning rates showed the opposite pattern from that predicted by the cue-choice congruency model (**Fig. 4k, left**). In the incongruent-manipulation group, however, the best-fitting model still incorporated cue-choice congruency-dependent learning, combined with differential cue or choice weighting during decision making. Moreover, the estimated learning rates qualitatively preserved the cue-choice congruency pattern observed without manipulation. (**Fig. 4l, left**).

Reorganizing the learning rates according to whether cue groups contained salient or non-salient cues (see **Methods**) revealed a shared pattern across both manipulation conditions. On rewarded trials, the cue group containing only non-salient cues tended to undergo stronger potentiation than the cue group containing the salient cue. On unrewarded trials, however, the cue group containing the salient cue tended to undergo stronger depression (**Fig. 4k,l right**). Interestingly, this learning-rate asymmetry may also help maintain greater dissociation between salient and non-salient cues by offsetting the additional influence that the manipulated cue gains through increased saliency. Together, these results support our hypothesis that neural activation elicited by luminance increase and subsequent fixation during feedback, whether driven by value-guided or exogenous attention, affects learning rates.

## Discussion

Here, using a multi-cue probabilistic learning task together with computational modeling, eye tracking, and saliency manipulation, we tested whether the brain solves credit assignment and adaptive learning jointly. Specifically, we asked whether cognitive processes involved in credit assignment shape learning rates to enhance discrimination between competing predictive cues. Our findings reveal a novel form of learning-rate asymmetry based on cue-choice congruency that is tightly linked to activation patterns shaped by value-guided overt attention during feedback. This asymmetry can be manipulated through cue saliency, which alters the activation and processing of cues. Specifically, on rewarded trials, learning rates were higher for cues that supported the chosen option, whereas on unrewarded trials, learning rates were higher for cues that opposed the chosen option. The cue-choice congruency effect was predicted by cue-choice congruency in gaze during feedback, indicating that learning depended on which cues were preferentially processed as reward outcomes were revealed. Cue-choice congruency-dependent learning, in turn, enhanced discrimination between predictive cues and supported more robust learning. Finally, manipulating cue activation and gaze through transient luminance increases during feedback altered the learning-rate asymmetry in the manner predicted by our hypothesis.

Thus, we suggest that asymmetric learning rates may arise from reward-dependent prioritization of choice-congruent cues and suppression of incongruent cues during value updating. Moreover, learning depends on which representations are active when reward outcomes are received, and even transient activation induced by external factors can influence updating, consistent with reward-dependent Hebbian learning (Frémaux & Gerstner, 2016; Gerstner et al., 2018). While low-level synaptic mechanisms such as metaplasticity can also produce such asymmetries (Farashahi, Donahue, et al., 2017; Khorsand & Soltani, 2017), our results point to a role for higher-level processes such as value-guided overt attention that can influence coactivation of task-relevant representations (Niv, 2019). Critically, overt attention may provide a mechanism for binding otherwise distinct representations, as fixation has been shown to align the encoding of option value in orbitofrontal cortex with the encoding of spatial location in lateral prefrontal cortex, causing both attributes of the attended option to be represented simultaneously despite their functional dissociation (Munet & Wallis, 2026).

Making learning rates depend on cue-choice congruency (the cue-choice congruency effect) helps address a key challenge in environments with multiple predictive cues. When cues provide conflicting information about outcomes, they generate prediction errors of different magnitudes. On rewarded trials, cues supporting the chosen option (congruent cues) already predict the outcome well and thus generate smaller prediction errors, whereas opposing cues (incongruent cues) generate larger prediction errors. If all cues are updated with the same learning rate, incongruent cues will change more, reducing the difference in predictive power between cues. On unrewarded trials, the opposite occurs: congruent cues generate larger negative prediction errors and are updated more, again reducing these differences. Although updates in both cases bias behavior toward the more rewarding option, equal learning rates diminish the discriminability of cues. Cue-choice congruency-dependent learning mitigates this effect by differentially updating cues according to their alignment with the choice, thereby preserving their relative predictive value. Moreover, this effect stabilizes cues linked to the more common outcome against expected uncertainty in reward, while keeping cues linked to the less common outcome sensitive to negative feedback and changes in reward associations.

Beyond identifying a mechanism that links learning-rate asymmetries to credit assignment, our results also provide a unified account of previously observed choice-dependent asymmetries in RL. Specifically, learning rates are typically higher for rewarded than unrewarded outcomes for the chosen option, but larger for unrewarded than rewarded outcomes for the unchosen option (Ciranka et al., 2022; Palminteri et al., 2017). We argue that this phenomenon, referred to as choice-confirmation bias, serves a similar functional role by preserving or amplifying value differences between the two options. Because the chosen option typically has higher value, equal learning rates reduce (stimulus or action) value differences when outcomes are similar across options (when the outcomes are different, value updates are in opposite direction). In contrast, asymmetric learning rates, which favor the chosen option on rewarded trials and the unchosen option on unrewarded trials, maintain or enhance this separation. As a result, these asymmetries improve robustness to noise in reward outcomes (Lefebvre et al., 2022; Palminteri et al., 2017). This increased robustness, however, comes at the cost of slower adaptation to environmental changes, reflecting an adaptability-precision trade-off (Farashahi, Rowe, et al., 2017; Khorsand & Soltani, 2017) and highlighting the need for additional mechanisms, such as change detection (Farashahi, Rowe, et al., 2017; Iigaya, 2016; Jang et al., 2015; McGuire et al., 2014; Piray & Daw, 2024). Interestingly, belief-dependent asymmetries in learning rates, where belief-consistent prediction errors are weighted more strongly than inconsistent ones, also help maintain differences in the predictive values of otherwise uninformative cues (Yazdanpanah et al., 2026). This advantage, however, comes with a downside: it can sustain biased perceptions driven by mere social suggestion.

Thus, we suggest that asymmetries in learning rates extend beyond cues, choices, and beliefs, reflecting a general principle by which the brain preserves discrimination among predictive cues. Although this can reduce flexibility or sustain socially driven biases when only a single cue, or no cue, is available, cue-choice congruency mitigates these costs by enabling greater precision for higher-prior cues and greater flexibility for lower-prior cues.

By separating the cues that drive choice from the options themselves, we were able not only to dissociate cue-based prediction processes from decision making, but also to distinguish how overt attention affects learning versus decision making. Specifically, gaze during the choice epoch was primarily biased toward higher-prior options, whereas during the feedback epoch it was driven by cue-choice congruency. While prior work has linked gaze to choice during deliberation (Krajbich et al., 2010; Smith & Krajbich, 2019) and to feedback processing during learning (Arbel et al., 2020), our paradigm enabled a direct comparison of these two phases within the same task, revealing distinct gaze patterns during decision making and feedback. Accordingly, variability in cue processing during choice did not predict the weights those cues exerted on decisions, suggesting that attentional influences on decision making are distinct from those on learning captured by the cue-choice congruency model (Leong et al., 2017; McGinty, 2019; Wang & Soltani, 2025). Our findings are consistent with the idea that learning prioritizes information that reduces uncertainty about the validity of the decision process itself, rather than about options in isolation (Bruckner et al., 2025).

Our results suggest that learning asymmetries can be driven by value-guided attention, a process naturally engaged to solve the credit assignment problem. This links credit assignment to adaptive learning-rate control, promoting more robust learning, and highlights the central role of attention in learning. It would be interesting to explore whether including such coupling can enhance learning in artificial systems (Neftci & Averbeck, 2019). Although our results are more consistent with value-guided attention modulating activity during feedback (Leong et al., 2017), choice-related activity may also generate eligibility traces that interact with subsequent reward signals, producing larger learning rates for congruent than incongruent cues on rewarded trials. This interpretation is consistent with attention-gated reinforcement learning frameworks, in which selective attention determines which representations are eligible for reward-modulated plasticity (Pozzi et al., 2020; Roelfsema & Ooyen, 2005; Vartak et al., 2017). Future work should determine neural mechanisms by which credit assignment alters learning rates, for example via eligibility traces (Izhikevich, 2007) or attention-dependent neuromodulation (Brzosko et al., 2019; Frémaux & Gerstner, 2016).

## Methods

### Experimental procedure

#### Participants

A total of 116 participants (40 males) were recruited from the Dartmouth College community, of whom 83 performed the task under unequal priors and 33 under equal priors. Of the 83 participants in the unequal-prior condition, ten were excluded from analysis (seven did not complete the task, two exhibited extremely biased choice behavior, and one reported inconsistent posterior estimates), leaving 73 participants (22 males; aged 17-33 years), including 37 with p(windy) = 0.75 and 36 with p(windy) = 0.25. A subset of these participants (N = 37; 19 from the p(windy) = 0.75 group and 18 from the p(windy) = 0.25 group) completed an additional saliency-manipulation session in which the saliency of one cue was transiently increased during feedback. Of the 33 participants in the equal-prior (control) condition (14 males; aged 19-39 years), one was excluded for not completing the task. All participants were naïve to the purpose of the study.

All experimental procedures were approved by the Dartmouth College Institutional Review Board and informed consent was obtained from all participants before the experiment. Participants were compensated with either monetary payment or T-points (extra credit points for courses in Dartmouth College’s Department of Psychological and Brain Sciences). Base compensation was $10 per hour or one T-point per hour, with performance-based bonuses of up to an additional $10 per hour.

#### Stimuli and choice options

The cues consisted of 12 unique grayscale geometric shapes, and the choice options were two grayscale weather predictions: windy and rainy. The cues and choice options were controlled for luminance contrast. The choice options were also matched in pixel count (approximately 1,700 pixels each) to prevent stimulus-driven attentional biases toward specific cues or choice options.

For each participant, four shapes were randomly selected for the main task and four for the practice session. The remaining four shapes were used in the saliency-manipulation session for the subset of participants who also performed the saliency-manipulation session. Each shape was randomly assigned one of four log-likelihood ratios (logLRs) favoring the windy option:

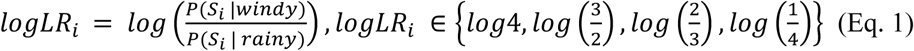

where *P*(*S_i_* | *windy*) represents the likelihood that cue *S_i_* was presented given that the windy option was the correct response. We label cues assigned with these *logLR* values as S1 to S4: S1 and S4 provide the strongest evidence in opposite directions and S2 and S3 provide weaker evidence in opposite directions.

#### Task

The main task included four blocks of choice trials and four blocks of estimation trials, alternating such that each choice block was followed by an estimation block. Each choice block contained 53 trials; the first three estimation blocks contained 8 trials each, and the final estimation block contained 53 trials. Prior to the main task, participants completed a practice session in which the prior probability of the windy option was set to 0.5 (p(windy)=0.5). The session consisted of one block of 15 choice trials and one block of 15 estimation trials.

Each choice trial began with a white fixation dot at the center of the screen, which turned yellow to signal the start of the trial. Participants were required to maintain fixation on the yellow dot for 1000 ms; otherwise, the trial did not begin. Once initiated, four cues were presented, and participants chose between two weather prediction options (windy vs. rainy) by pressing ‘F’ or ‘J’ to select the left or right option, respectively. We refer to this as the choice epoch. After a response was made, a circle highlighted the chosen option, followed by reward feedback (correct/incorrect) and an update to a reward bar indicating the recent accumulation of points. We refer to this as the feedback epoch. When the reward bar reached 10 points, it turned gold and reset to one point on the next rewarded trial. Participants were instructed to collect as many gold bars as possible, as these were converted into monetary rewards at the end of the experiment.

During each estimation trial, participants reported the probability that the windy or rainy option would be rewarded given a single cue or a combination of cues. The first three estimation blocks consisted of one-cue trials (four for windy and four for rainy). The final block included 8 one-cue trials, followed by 10 unique two-cue trials and 35 unique four-cue trials.

The control task followed the same structure, except that it included only three blocks of each type (choice and estimation), and participants provided single-cue estimations only. Eye tracking was not performed during the control experiment.

#### Eye tracking

Eye-tracking data was acquired using an EyeLink 1000 Plus system with a sampling rate of 500 Hz. Before the task began, a 9-point grid calibration was performed for each participant. During choice blocks, trial onset was gaze-contingent, requiring participants to fixate within ±50 pixels of the gray circle at the center of the screen for 1000 ms before each trial began. Once a trial started, participants were free to look anywhere on the screen.

#### Cue saliency manipulation

A subset of participants (N = 37) completed an additional session in which the luminance of one of the two least informative cues (S2 or S3) was transiently increased twice at the onset of reward feedback, causing the cue to briefly blink and appear more salient. This within-subject design allowed us to compare each participant’s behavior with and without the manipulation.

The saliency-manipulation session followed the same structure as the main task, except that whenever a predefined cue (S2 or S3) appeared, its luminance was transiently increased at feedback onset to draw exogenous attention to that cue. The identity of the salient cue was fixed for each participant throughout the experiment and the manipulation was restricted to the feedback epoch. To dissociate exogenous attentional capture from value-guided attention, the salient cue was chosen such that for 19 participants it favored the higher-prior option (S3 if p(windy) = 0.25; S2 if p(windy) = 0.75), whereas for 18 participants it favored the lower-prior option (S2 if p(windy) = 0.25; S3 if p(windy) = 0.75). For some analyses, participants were divided into congruent-and incongruent-saliency groups based on the saliency-congruency index (see Saliency-congruency index).

To combine data across the two prior-probability groups, we re-labeled cues according to their predictive value for the higher-prior option in each condition. For participants with p(windy) = 0.75, the higher-prior option was windy and the original cue labels were retained (S1′–S4′ = S1–S4). For participants with p(windy) = 0.25, the higher-prior option was rainy and the cue labels were reversed (S1′ = S4, S2′ = S3, S3′ = S2, S4′ = S1). Under this convention, S1′–S4′ always denotes cues ordered from most to least predictive of the higher-prior option. We use this notation throughout the paper whenever analyses are pooled across prior-probability groups.

### Data Analysis

#### Behavioral analysis

For a given trial, the log posterior odds that reward is assigned to the windy rather than the rainy option is given by:

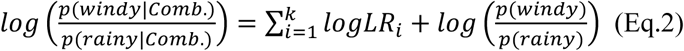

where *p*(*windy*|*Comb*.) and *p*(*rainy*|*Comb*.) denote the posterior probabilities that the windy and rainy options, respectively, are rewarded (correct option) given the presented combination of *k* shapes, and *logLR_i_* is the log-likelihood ratio associated with cue *S_i_*. Performance was quantified as the proportion of trials on which participants chose the option with the higher posterior probability of reward, as determined by Eq. 2.

To characterize the temporal dynamics of learning, we fit the time course of performance with an exponential function

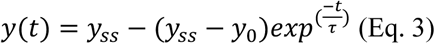

where *y*_0_ represents the initial performance, *y_ss_* is the steady state performance, *τ* is the time constant for approaching the steady state, and *t* represents the trial number.

To quantify the influence of individual cues on choice behavior, we fit a logistic regression model of the form

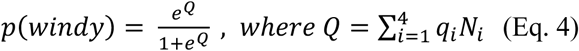

where *N_i_* is the number of appearances of cue *i* in the presented combination and *q_i_* is the corresponding regression coefficient. These coefficients, referred to as subjective weights of evidence (SWOEs), quantify the contribution of each cue to participants’ choices. To examine how cue weighting evolved over learning, SWOEs were estimated separately for each participant using a sliding window of 45 consecutive trials advanced one trial at a time.

#### Cue-congruency index

The cue-choice congruency (CC) index is defined as the average of two differences: (1) the learning rate for cues congruent minus incongruent with the chosen option on rewarded trials, and (2) the learning rate for cues incongruent minus congruent with the chosen option on unrewarded trials. This average is then normalized by the inverse temperature parameter (1/β; Eq. 5) to account for inter-participant variability in sensitivity to choice. A higher CC index indicates stronger asymmetric learning, with greater updating of congruent cues on rewarded trials and incongruent cues on unrewarded trials.

#### Saliency-congruency index

For each participant, we computed a saliency-congruency index, defined as the log ratio between the number of trials on which the salient cue was congruent with the participant’s choice and the number of trials on which it was incongruent with the choice. For some analyses, participants with a positive index were assigned to the congruent-saliency group, whereas those with a negative index were assigned to the incongruent-saliency group.

#### Eye data analysis

For each cue and choice option, we defined an area of interest extending 10 pixels beyond its boundary. Raw gaze samples (500 Hz) were grouped into fixations using a spatial and temporal criterion: consecutive samples lasting at least 100 ms and remaining within 25 pixels (<1° visual angle) of the running centroid were classified as a fixation. Samples not belonging to a qualifying fixation were excluded from dwell-time analyses.

For each trial, dwell time for a cue was computed as the total fixation time across all occurrences of that cue, normalized by the number of occurrences and expressed as a proportion of trial duration. Dwell time for each choice option was computed as the proportion of trial duration spent fixating that option. The reported proportion of dwell time on a given cue or choice option was then calculated relative to the total fixation time on all task-relevant elements (all cues and choice options) and averaged across trials.

Eye-movement analyses were performed separately for the choice and feedback epochs. The choice epoch extended from the end of the fixation period to the participant’s response, whereas the feedback epoch extended from feedback onset (“correct” or “incorrect”) to the end of the trial.

#### Regression analysis of congruency-dependent gaze during feedback

To test whether gaze during the feedback epoch was jointly modulated by trial outcome and cue–choice congruency, we fit linear mixed-effects models using MATLAB’s *fitlme* function. All models included subject-specific random intercepts to account for repeated measurements and individual differences in baseline gaze. Models with random slopes for congruency, outcome, or both were compared using log-likelihood, with the best-fitting model including random slopes for both predictors. Fixed-effect estimates, standard errors, t-statistics, and p-values were extracted from the fitted model.

#### Gaze-based prediction of choice

To examine how dwell time during the choice epoch influenced decisions, we fit participants’ trial-by-trial choices using logistic regression models implemented with MATLAB’s *fitglm* function. Specifically, two gaze predictors were computed for each trial: (1) differential dwell time on the cue groups, defined as the difference in dwell time between S1+S2 and S3+S4; and (2) differential dwell time on the choice options, defined as the difference in the dwell time between the windy and rainy options. Trials with missing values for either predictor or choice were excluded. To place participants on a common scale, choices and gaze predictors were sign-flipped for participants with p(windy) = 0.25, such that a choice value of 1 always indicated selection of the higher-reward option and positive gaze values indicated greater attention toward cues or options favoring that option.

Predictive accuracy was assessed using within-subject 5-fold cross-validation. For each participant, models were trained on four folds and tested on the held-out fold, with predicted probabilities thresholded at 0.5. Accuracies were averaged across folds to obtain a single value per participant and model. Models were compared at the group level using paired t-tests, and Cohen’s d was computed from the paired differences. Each gaze model was compared against a null model, and the combined model was additionally compared with the option-gaze model to assess whether cue gaze provided predictive information beyond option gaze alone.

#### Effect of cue–choice congruency asymmetry on performance and sensitivity to evidence

To assess whether individual differences in cue–choice congruency (CC) asymmetry were associated with behavior, we computed two measures separately for an early phase (first 70 trials) and a late phase (final 70 trials): (1) task performance, defined as the proportion of correct choices; and (2) sensitivity to objective evidence, quantified as the Spearman correlation between the total evidence provided by the presented cues and participants’ choices. For each measure and phase, we fit a generalized linear model with CC group as a categorical predictor to test whether behavior differed across groups.

### Computational models

#### Models descriptions

Our models build on the neural circuit model of probabilistic inference proposed by Soltani and Wang (Soltani & Wang, 2010), in which cue–outcome associations are encoded by plastic synapses that undergo reward-modulated stochastic Hebbian learning. We extend this framework by assuming that additional cognitive processes, such as attention, can influence the plasticity of these synapses (learning rates) through neuromodulatory mechanisms, thereby shaping the learning of cue-outcome associations.

All models share a common neural architecture consisting of three interconnected circuits. The first is a cue-encoding circuit composed of sensory neurons selective for individual shapes/cues. These neurons are assumed to reside in visual areas, particularly the inferotemporal cortex, where shape-selective neurons have been extensively documented (Tanaka, 1996; Zhivago & Arun, 2016). This circuit encodes sensory cues and projects to the value-encoding circuit through plastic synapses that mediate learning.

The value-encoding circuit receives inputs from the cue-encoding circuit through synapses that undergo Hebbian plasticity, modulated by reward and other cognitive processes. It consists of neurons that encode the reward values associated with the two choice options (rainy and windy) and is assumed to reside in regions such as the basal ganglia, anterior cingulate cortex, or dorsolateral prefrontal cortex, where value-coding neurons have been identified (Kim et al., 2008; Rushworth & Behrens, 2008; Samejima et al., 2005). We further assume that the activity of these neurons can be gain-modulated by cognitive processes such as attention, leading to stronger responses to selected inputs from the cue-encoding circuit.

The decision-making circuit integrates value information from the value-encoding circuit to generate choices. It consists of two competing neural populations, each representing one of the two choice options (rainy and windy). The relative activity of these populations determines the probability of selecting a given option. Specifically, the probability of choosing the windy option on trial *t* is given by

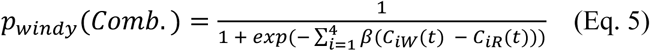

where *C_i_*_W_(*t*) and *C_iR_*(*t*) denote the strength of synapses from neurons encoding cue *i* onto neurons encoding the value of the windy and rainy options, respectively, summed across all instances of cue *i* in the presented combination, and β controls the sensitivity of the decision circuit to differences in the values of the two options. We refer to *C_i_*_W_(*t*) and *C_iR_*(*t*) as synaptic strengths; together, they provide cue-specific estimates of the predictive relationship between each cue and the two outcomes, and thus of the posterior belief associated with that cue.

The selected action is then executed, resulting in either reward or no reward. Reward outcomes modulate synaptic plasticity such that rewarded trials induce potentiation of plastic synapses, whereas unrewarded trials induce synaptic depression. Specifically, following a rewarded trial, the synaptic strengths from the presented cues onto value-encoding neurons selective for the chosen option (here, the windy option) are updated according to:

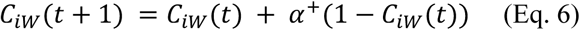

Where α^+^ is the potentiation rate or the learning rate for rewarded trials. On non-rewarding trials, the synaptic strengths are updated as:

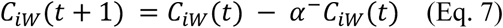

where α^−^ is the depression rate or the learning rate for unrewarded trials. All other synapses remain unchanged, including synapses from neurons encoding cues that were not presented and synapses from neurons encoding presented cues onto value-encoding neurons selective for the unchosen option.

Overall, we tested 10 candidate models. In addition to a baseline model with a single learning rate, we considered three forms of learning-rate modulation, in which learning rates depended on reward outcome, cue category, or cue–choice congruency. Each of these learning models was combined with one of three decision-making schemes: no additional weighting, differential cue weighting, or differential option weighting (see below).

### Model 1: Single learning rate

This baseline model assumes symmetric learning, with cue-outcome associations updated using the same learning rate on rewarded and unrewarded trials. Consequently, positive and negative prediction errors modify synaptic strengths at equal rates.

### Model 2: Separate learning rates for rewarded and unrewarded outcomes

This model allows learning to differ depending on the outcome of the trial by assigning separate learning rates to rewarded and unrewarded outcomes (separate α^+^ and α^−^).

### Model 3: separate reward-dependent learning rates for different cue groups

This model assumes that learning rates depend on both reward outcome and cue category. Specifically, cues were divided into two groups according to the direction of the evidence they provide: cues favoring the higher-prior option and cues favoring the lower-prior option. We hypothesized that these cue groups may be updated at different rates, potentially due to differential processing or attentional allocation. Specifically, following a rewarded trial, the synaptic strengths from the presented cues onto value-encoding neurons selective for the chosen option (here, the windy option) are updated according to:

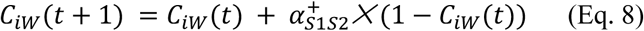

where 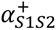 is the potentiation rate of synapses from cues S1 and S2. Concurrently, the synaptic strengths of presented cues carrying evidence against the windy option are updated according to:

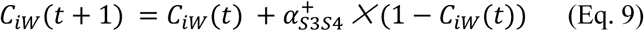

where 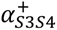 is the potentiation rate of synapses from cues S3 and S4. Following an unrewarded trial in which the windy option was chosen, synaptic strengths from presented cues carrying positive (Eq. 10) or negative (Eq. 11) evidence onto value-encoding neurons selective for the windy option are updated according to:

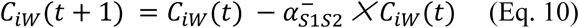

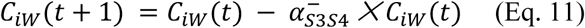

where 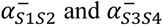 are the depression rates of synapses from cues S1 and S2, and S3 and S4, respectively.

### Model 4: cue-choice congruency-dependent learning rates

This model assumes that learning rates depend on whether a cue is congruent or incongruent with the chosen option. Specifically, cues supporting the chosen option and cues opposing it are updated at different rates. Separate potentiation and depression rates are assigned to congruent and incongruent cues 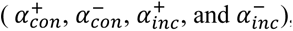, such that synaptic updates depend jointly on trial outcome (rewarded or unrewarded) and cue-choice congruency.

In addition to asymmetries in learning, we considered the possibility that inputs are differentially weighted during decision making. We therefore implemented two decision-making schemes: option weighting, in which inputs favoring the windy and rainy options are weighted separately, and cue weighting, in which cues favoring the higher-and lower-prior options are weighted separately.

Models 5 and 6 combine the learning rule of Model 2 (separate learning rates for rewarded and unrewarded outcomes) with option weighting and cue weighting, respectively. In the option-weighting variant, separate parameters (β_windy_ and β_rainy_) scale inputs favoring each choice option:

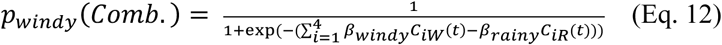

In the cue-weighting variant, separate parameters (β_S1S2_ and β_S3S4_) scale the contributions of the two cue groups:

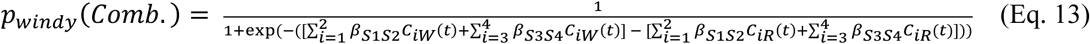

The same two decision-making schemes were then combined with the learning rules of Models 3 and 4, yielding four additional models. Specifically, Models 7 and 8 combine cue-category-dependent learning (Model 3) with option weighting and cue weighting, respectively, whereas Models 9 and 10 combine cue-choice congruency-dependent learning (Model 4) with option weighting and cue weighting, respectively.

### Mapping estimated learning rates onto salient and non-salient cues

To determine whether the models that best explained behavior during the saliency-manipulation session assigned different learning rates to cue groups containing the salient versus non-salient cues, we reorganized the estimated learning-rate parameters accordingly. Because the identity of the salient cue depended on both manipulation condition and prior-probability condition, the specific cue group designated as “salient” differed across participants but was known for each participant and used to relabel learning-rate parameters before pooling.

For the cue-prior model, which estimates separate potentiation (α^+^) and depression (α^−^) rates for the two cue groups (S1S2 and S3S4), we relabeled the estimated learning rates according to whether the corresponding cue group contained the salient cue or only non-salient cues and then pooled data across prior-probability conditions. This procedure yielded estimates of potentiation and depression rates for salient and non-salient cue groups separately within each saliency-manipulation condition.

For the cue–choice congruency model, which estimates separate learning rates for congruent (α^+^_con_, α^−^_con_) and incongruent (α^+^_inc_, α^−^_inc_) cues, we mapped learning rates onto salient and non-salient cue groups according to the dominant relationship between saliency and choice. In the congruent-saliency condition, where the salient cue was more often congruent with the chosen option, congruent learning rates were assigned to the salient and aligned cue group, whereas incongruent learning rates were assigned to the non-salient cue group. In the incongruent-saliency condition, where the salient cue was more often incongruent with the chosen option, this assignment was reversed. Because cue–choice congruency varies from trial to trial whereas saliency is fixed at the cue-group level, this mapping is approximate and reflects the predominant alignment between saliency and choice. For both models, group means and standard errors were computed for the resulting potentiation (α^+^) and depression (α^−^) rates of the salient and aligned cue group and the non-salient cue group.

### Model fitting and model selection

Model parameters were fit separately to each participant’s trial-by-trial choices by maximizing the likelihood of the observed choices under each model. Parameter optimization was performed in MATLAB using *fminunc* to minimize the negative log-likelihood of the predicted choice probabilities. To reduce the risk of local minima, the fitting procedure was repeated 50 times per participant with different initial parameter values, and the best-fitting solution was retained.

To compare models while accounting for differences in model complexity, we used Bayesian model selection (BMS; Rigoux et al., 2014; Stephan et al., 2009) based on the Bayesian information criterion (BIC), which served as an approximation to model evidence. For each model, we report the mean BIC, the estimated frequency (EF; the posterior estimate of the proportion of participants best described by the model), and the protected exceedance probability (PXP; the probability that a model is more likely than any competing model after accounting for chance differences in model frequencies).

The Bayesian omnibus risk, which quantifies the probability that observed differences in model frequencies arose by chance, was 4.98 × 10⁻^18^, indicating strong evidence that the models differed in their ability to explain participants’ behavior.

### Model and parameter recovery

To assess the reliability of model comparison and parameter estimation, we conducted model recovery and parameter recovery analyses. For each model, synthetic datasets were generated using parameters sampled from empirical distributions derived from the fitted participant data. To construct these distributions, we first identified and excluded outliers in the sensitivity parameter (β) using the 1.5 × IQR criterion. Kernel density estimates were then fit to the remaining parameter values, and synthetic parameters were sampled from these distributions. This procedure generated parameter values that closely matched those observed in participants without imposing a specific parametric form.

Using the sampled parameters, we simulated 75 datasets of 212 trials for each of the 10 candidate models. The simulated choices were then fitted with all 10 models and compared using BMS. Model recovery was assessed using the EF and PXP, and the true generating model was correctly identified in all cases. Parameter recovery was evaluated by calculating the Pearson correlation between the true and recovered parameter values.

Because the empirical parameter distributions were derived from fits of multiple models to the same behavioral dataset, this retrospective validation procedure is conservative and may underestimate the true discriminability among models.

## Acknowledgements

We are grateful to Jamie Lim and Seo Yoon Park for assistance with data collection, and to Aryan Yazdanpanah for discussions and for reviewing portions of the analyses and associated code. We also thank Daeyeol Lee for helpful comments on the manuscript. This work was supported by National Science Foundation CAREER Award (BCS1943767) to A.S.

---

**Figure S1.**
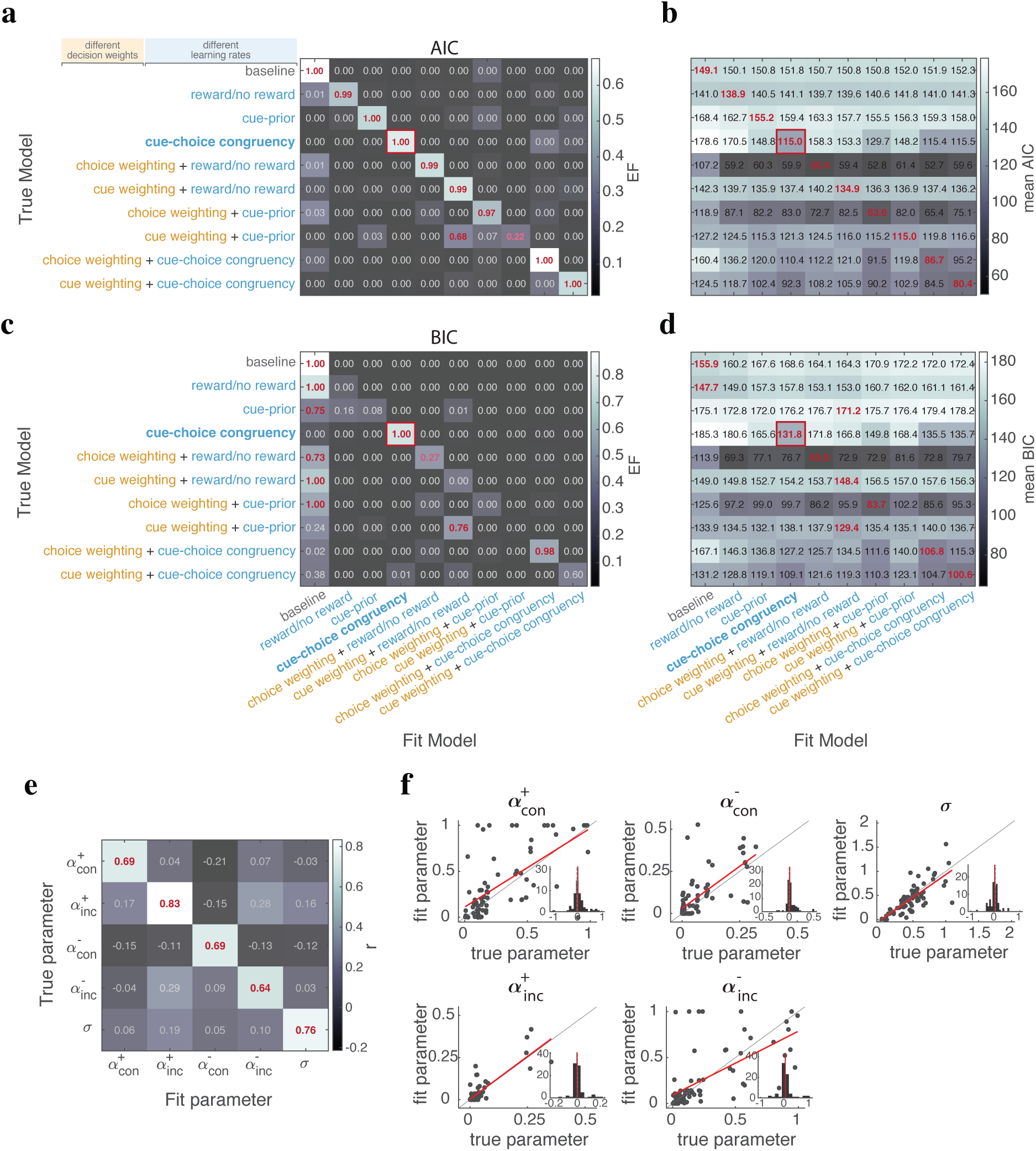
Model and parameter recovery analyses. (**a–d**) Model recovery results for the 10 candidate models. Simulated datasets were generated from each model using parameters sampled from empirical parameter distributions and then fit with all candidate models. Panels (a) and (c) show Bayesian model selection results using AIC and BIC, respectively, as approximations of model evidence. Protected exceedance probability (PXP) is shown in each cell, and estimated frequency (EF) is indicated by cell color. Panels (b) and (d) show the corresponding mean AIC and BIC values across simulated datasets. Higher EF and PXP values indicate stronger support for a model, whereas lower AIC and BIC values indicate a better fit. For each simulated model (rows), the best-fitting recovered model (columns) is indicated by a red square; cells with the highest PXP and lowest AIC/BIC values are highlighted in red. (**e,f**) Parameter recovery for the best-fitting model. Pearson correlation coefficients between true and recovered parameter values across simulations are shown by both cell color and text in panel e. Significant correlations after Bonferroni correction are highlighted in red. Panel (f) shows scatter plots comparing recovered and true parameter values for each model parameter. The red line shows the linear fit corresponding to the Pearson correlation coefficient. Insets show distributions of estimation errors (recovered minus true parameter values); dashed vertical lines indicate zero error and solid red lines indicate the median estimation error. Together, these analyses demonstrate reliable identification of the generating model and accurate recovery of model parameters.

**Figure S2.**
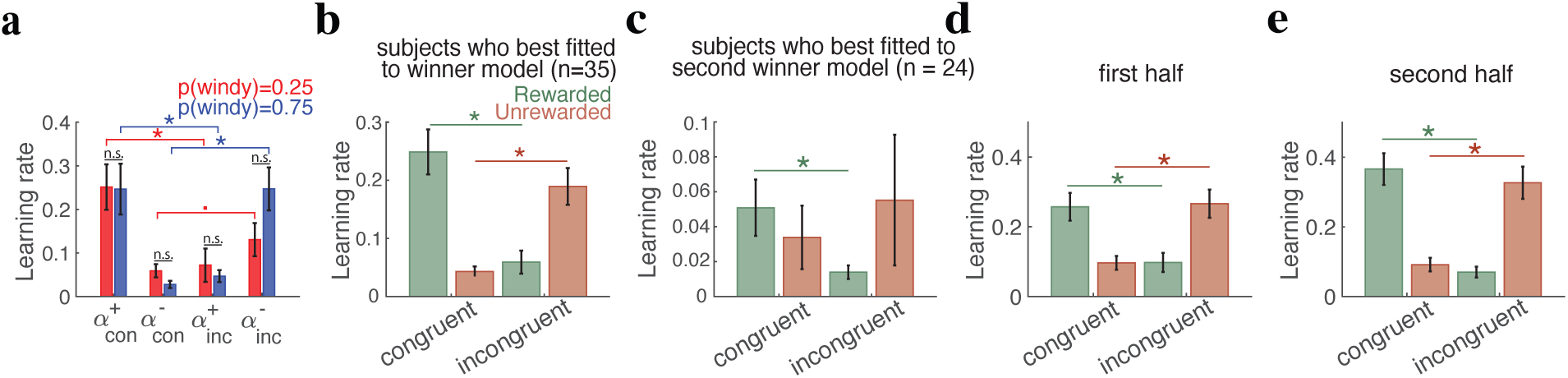
Robustness of cue-choice congruency-dependent learning across priors, participants, and time course of experiment. (**a**) Learning rates for congruent and incongruent cues on rewarded (α^+^_con_ and α^+^_inc_) and unrewarded (α^−^_con_ and α^−^_inc_) trials, shown separately for participants with prior probabilities of 0.25 and 0.75 (mean ± SEM). Asterisks indicate significant within-group differences between congruent and incongruent learning rates (Wilcoxon signed-rank test, p<0.05); filled circles (•) indicate marginal significance (p=0.05). Learning rates did not differ significantly between prior-probability groups for any parameter (Wilcoxon rank-sum test) (**b**) Estimated learning rates for participants best fit by the overall best-fitting model. (**c**) Estimated learning rates for participants best fit by the second best-fitting model (cue–choice congruency with cue weighting). (**d,e**) Learning rates estimated separately from the first (d) and second (e) halves of the experiment. The cue-choice congruency asymmetry was present in both the first half (Wilcoxon signed-rank test; rewarded: z(con−inc) = 3.96, p = 7.29 × 10^−5^, d = 0.46; unrewarded: z(inc−con) = 4.01, p = 6.20 × 10^−5^, d = 0.47) and second half of the experiment (rewarded: z(con−inc) = 5.72, p = 1.03 × 10^−8^, d = 0.67; unrewarded: z(inc−con) = 4.47, p = 7.54 × 10^−6^, d = 0.52). Direct comparison between halves showed that the cue–choice congruency effect was stronger in the second half of the experiment for rewarded trials (Wilcoxon signed-rank test: z=2.21, p=0.026, d=0.26) and marginally stronger for unrewarded trials (z=1.95, p=0.050, d=0.23), indicating that cue–choice congruency-dependent updating strengthens over the course of learning.

**Figure S3.**
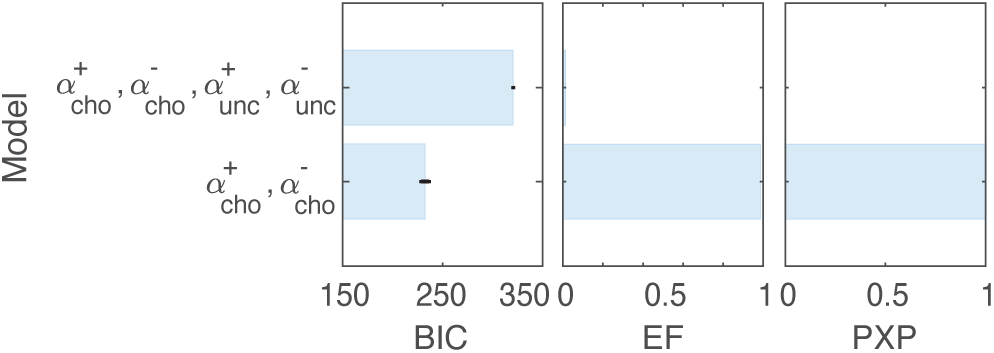
Separate learning rates for the unchosen option do not improve model fit. Comparison between a model with separate learning rates for rewarded and unrewarded outcomes of the chosen option (α^+/-^_cho_ ≠ α^+/-^_unc_) and an extended model that additionally included separate learning rates for updates to the unchosen option. Models were compared using BIC, estimated frequency (EF), and protected exceedance probability (PXP), where lower BIC and higher EF/PXP indicate better fit. Bayesian model selection favored the simpler model, indicating that learning asymmetries were specific to updates of the chosen option and that allowing separate learning rates for the unchosen option did not improve explanatory power.

**Figure S4.**
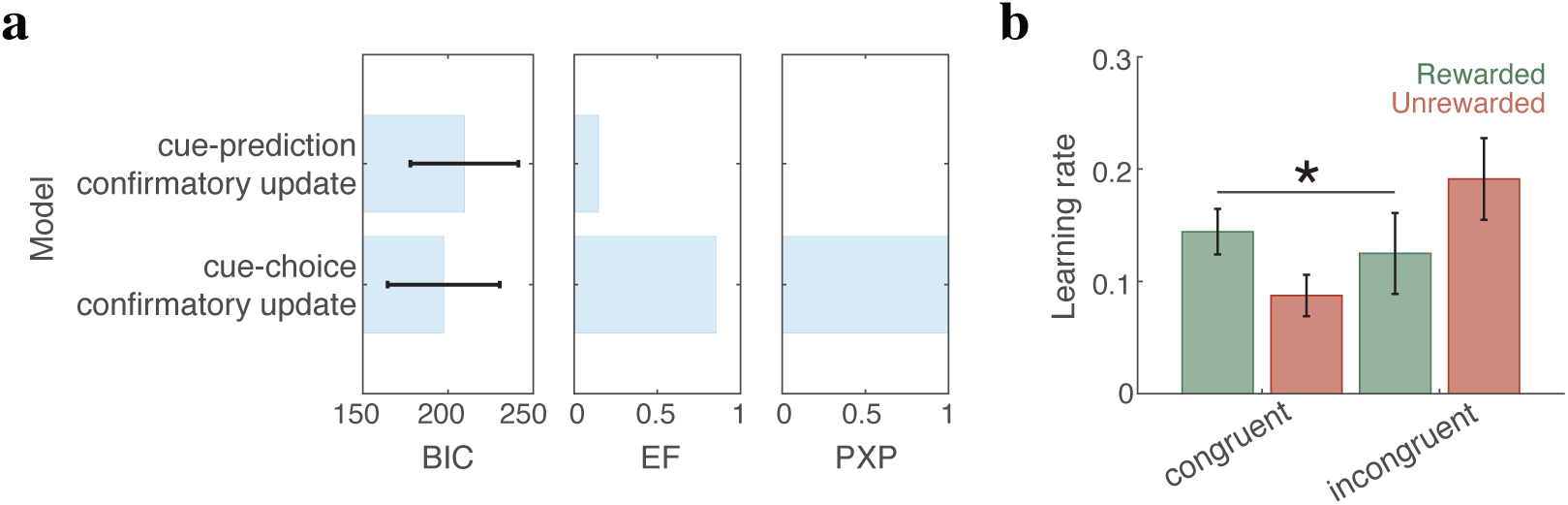
Choice behavior is better captured by cue–choice congruency than by cue–prediction congruency. Comparison of models in which learning-rate asymmetries depend either on the alignment between cues and the participant’s actual choice (cue–choice congruency model) or on the alignment between cues and the predicted option, defined as the option receiving greater overall input on a given trial (cue–prediction congruency model). (**a**) Model comparison using BIC, estimated frequency (EF), and protected exceedance probability (PXP). Lower BIC and higher EF/PXP indicate better model fit. Bayesian model selection favored the cue–choice congruency model over the cue–prediction congruency model. (**b**) Estimated learning rates for cues congruent and incongruent with the predicted option on rewarded and unrewarded trials. On rewarded trials, cues congruent with the predicted option showed stronger potentiation than incongruent cues (Wilcoxon signed-rank test: z=3.73, p=1.87×10^−4^, d=0.43), whereas no significant difference was observed on unrewarded trials (z=1.48, p=0.137, d=0.17). Together, these results indicate that learning asymmetries are more strongly tied to alignment with the participant’s actual choice than with internally generated predictions.

**Figure S5.**
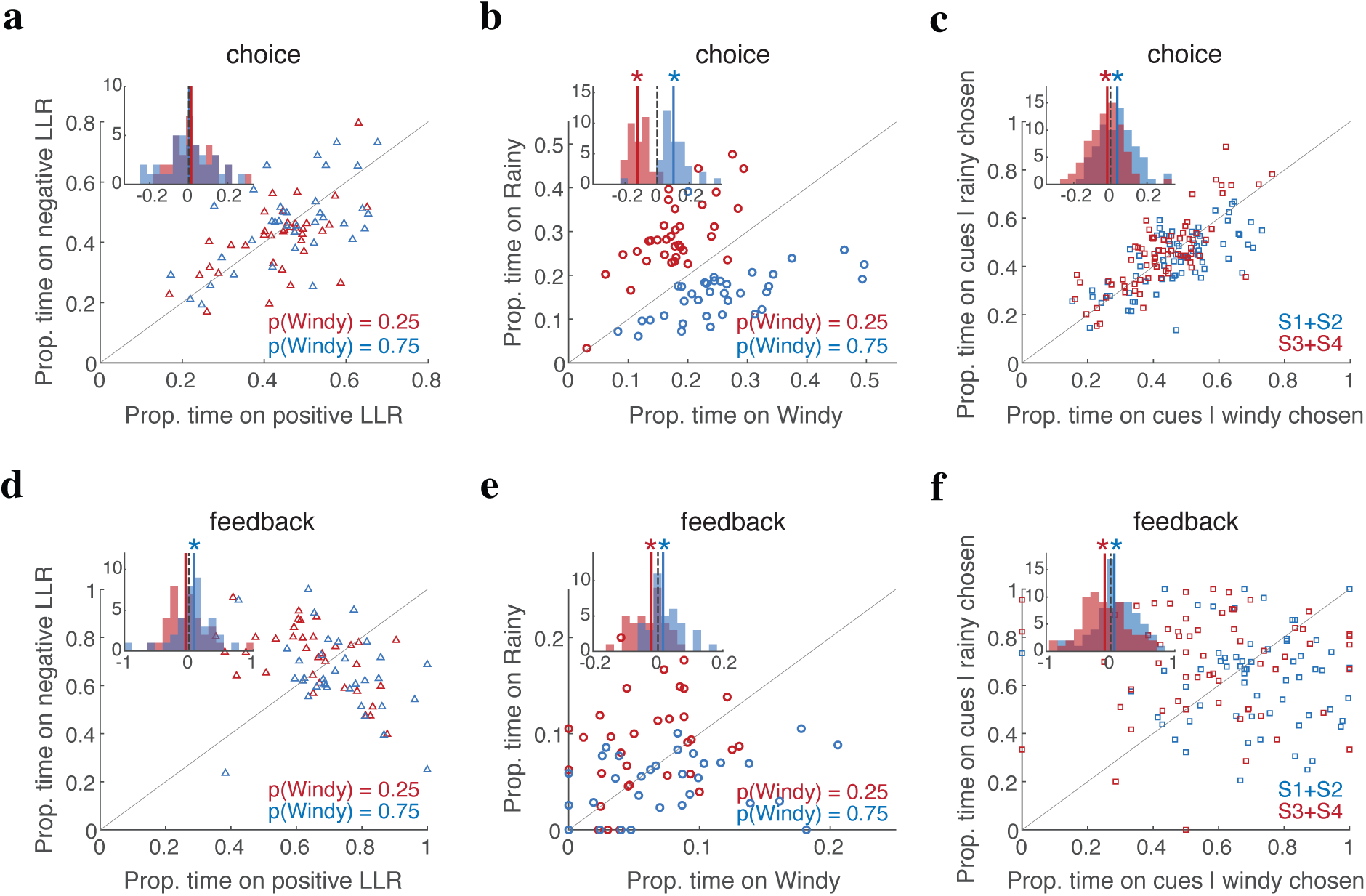
Cue-choice congruency is reflected in the pattern of dwell time. Dwell time on cues and choice options during the choice and feedback epochs. (**a**) Mean proportion of dwell time on cues with negative log-likelihood ratios (LLRs; supporting the rainy option) versus positive LLRs (supporting the windy option) during the choice epoch, expressed relative to the total dwell time on all task-relevant elements (cues and choice options), shown separately for the two priors. Histograms show the distribution of dwell-time differences between the two cue groups (S1+S2) -(S3+S4), with solid vertical lines indicating medians. (**b**) Mean proportion of dwell time on the rainy versus windy choice options during the choice epoch, shown separately by prior condition. Histograms show the difference in dwell time (windy−rainy); solid vertical lines indicate medians, and asterisks denote significant deviations from zero (Wilcoxon signed-rank test, p<0.05). Participants preferentially fixated on the higher-prior option. (**c**) Mean proportion of dwell time on cues congruent versus incongruent with the participant’s choice during the choice epoch. Histograms show the corresponding dwell-time differences, with medians indicated by solid vertical lines. Participants showed a bias toward cues supporting the chosen option. (**d–f**) Same analyses as in panels a–c, but during the feedback epoch. During feedback, participants tended to fixate on cues supporting the higher-prior option (d) and on higher-prior choice option itself (e). They also fixated cues congruent with their choice more than incongruent cues (f), consistent with the cue–choice congruency effect observed in learning rates.

**Figure S6.**
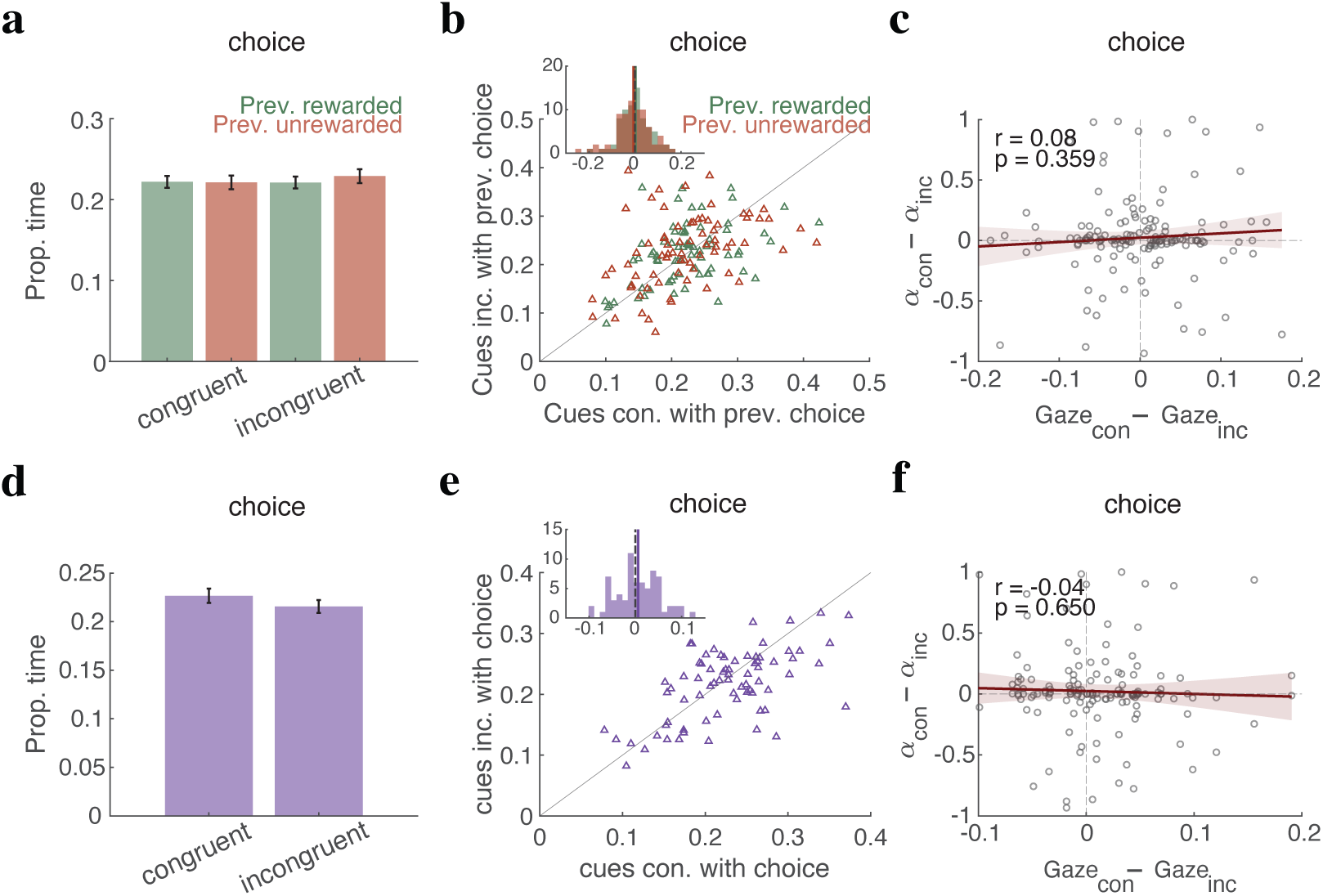
Patterns of gaze on subsequent trials and during choice epoch are not linked to learning-rate asymmetry. To test whether cue-choice-outcome interactions influenced subsequent decision making, we examined dwell time during the choice epoch for cues congruent or incongruent with the previous trial’s choice, separately for previously rewarded and unrewarded trials. (**a,b**) Mean proportion of dwell time on cues congruent versus incongruent with the previous choice, shown separately following rewarded and unrewarded trials. The inset histogram shows the distribution of dwell-time differences between congruent and incongruent cues. No significant differences in dwell time were observed following either rewarded (z=0.12, p=0.901) or unrewarded trials (z=−0.646, p=0.518; Wilcoxon signed-rank tests). (**c**) Relationship between the dwell-time difference for cues congruent versus incongruent with the previous choice and the corresponding learning-rate difference. The solid red line indicates the regression fit, with the shaded band showing the 95% confidence interval. (**d,e**) Mean proportion of dwell time on cues congruent versus incongruent with the chosen option during the choice epoch, with the inset histogram showing the distribution of dwell time differences. Participants did not show a significant dwell time bias toward congruent versus incongruent cues (z=1.37, p=0.16). (**f**) Relationship between the dwell time difference during the choice epoch and learning-rate asymmetry. Individual differences in dwell time did not predict learning-rate asymmetry (r=−0.04, p=0.65).

**Figure S7.**
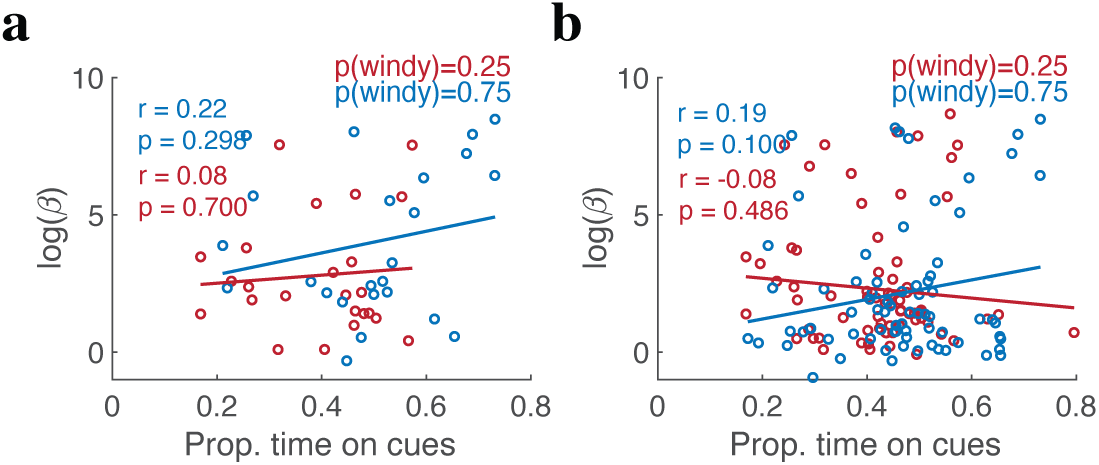
**Gaze patterns during the choice epoch do not predict cue weighting during decision making**. Relationship between cue-specific decision weights and dwell time during the choice epoch. (**a**) Log-transformed decision weights for each cue group (S1S2 vs. S3S4) plotted against the average dwell time allocated to that cue group for participants whose behavior was better fit by the model with separate cue-specific decision weights (N=24), shown separately for the p(windy)=0.25 and p(windy)=0.75 prior conditions. Solid lines indicate linear regression fits. No significant relationship was observed either overall (r=0.23, p=0.102) or within the individual prior groups (p(windy)=0.25: r=0.08, p=0.700; p(windy)=0.75: r=0.22, p=0.298). (**b**) Same analysis performed across all participants, again showing no significant relationship between cue weighting and dwell time (overall: r=0.24, p=0.103; p(windy)=0.25: r=−0.08, p=0.486; p(windy)=0.75: r=0.19, p=0.099). Together, these results indicate that dwell time during choice does not account for cue-weighting effects during decision making.

**Figure S8.**
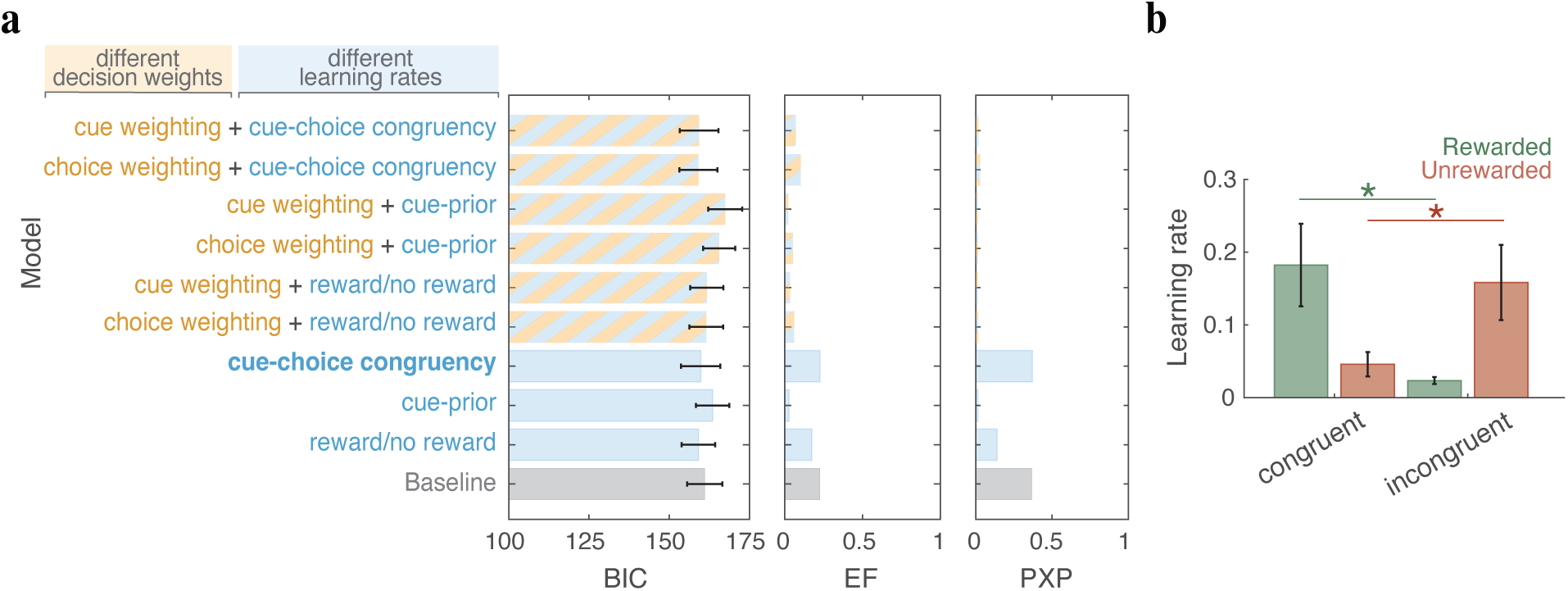
The cue-choice congruency effect is preserved under equal prior probabilities. (**a**) Model comparison for the control group with equal prior probabilities (N=32) using BIC, estimated frequency (EF), and protected exceedance probability (PXP). Lower BIC values indicate better model fit, whereas higher EF and PXP indicate stronger evidence for a model. The cue–choice congruency model (bold), which allows learning rates to differ for cues that support versus oppose the chosen option on rewarded and unrewarded trials, provided the best overall fit (PXP=0.37). The baseline model with a single learning rate and the model with separate learning rates for rewarded and unrewarded outcomes provided the second-and third-best fits, respectively (PXP=0.36 and PXP=0.14). (**b**) Estimated learning rates from the best-fitting model for cues congruent and incongruent with the chosen option, shown separately for rewarded and unrewarded trials. Error bars indicate SEM across participants, and asterisks denote significant differences (Wilcoxon signed-rank test, p<0.05). The cue–choice congruency effect remained significant under equal priors, with stronger updating of congruent than incongruent cues on rewarded trials (z(con−inc)=4.30, p=1.70×10^−5^, d=0.76) and stronger updating of incongruent than congruent cues on unrewarded trials (z(inc−con)=2.61, p=0.008, d=0.46).

**Figure S9.**
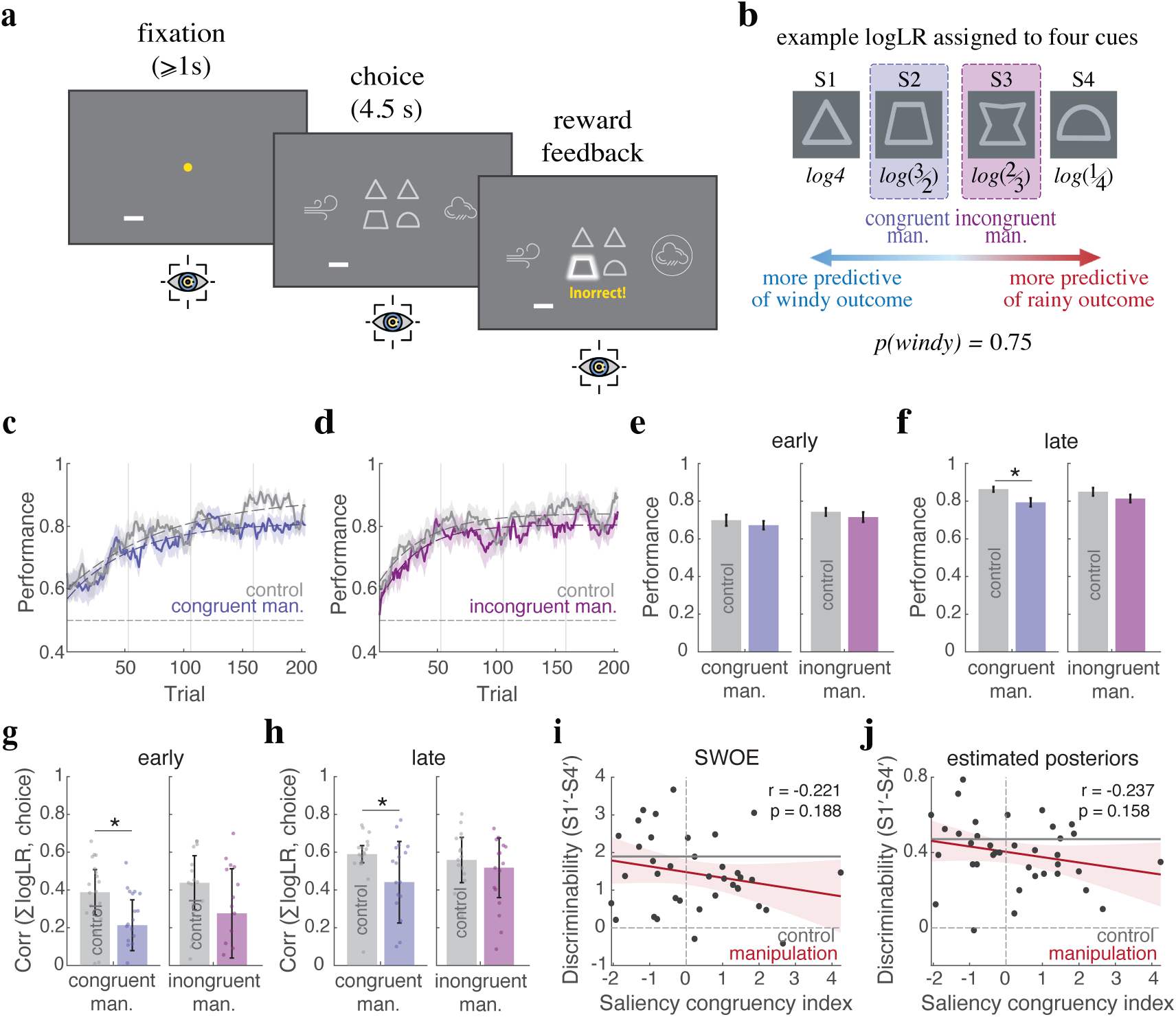
Saliency manipulation paradigm and disruption of cue-outcome associations. (**a**) Trial structure during saliency-manipulation sessions. At feedback onset (“correct” or “incorrect”), the luminance of a predefined cue was transiently increased to manipulate exogenous attention while gaze was continuously recorded. (**b**) Example cues and their evidence (logLR) for the windy option. Depending on prior condition (p(windy)=0.25 or 0.75), congruent manipulations increased the saliency of the cue aligned with the higher-prior option, whereas incongruent manipulations targeted the opposing cue. (**c,d**) Time course of performance for congruent (c) and incongruent (d) saliency-manipulation groups compared with their within-participant control sessions without manipulation (gray). Performance was defined as the proportion of trials on which the higher-posterior option was chosen. Shaded regions indicate SEM, vertical bars indicate estimation blocks, and dashed lines show exponential fits. (**e,f**) Average performance during the early (e; first 70 trials) and late (f; last 70 trials) phases of learning. Congruent saliency manipulation significantly reduced late-phase performance relative to control, whereas incongruent manipulation did not. Asterisks indicate significant differences (Wilcoxon signed-rank test, p<0.05). (**g,h**) Correlation between the summed objective weight of evidence (sum logLR) carried by the presented cues and participants’ choices during the early (g) and late (h) phases of the experiment. Congruent manipulation reduced the alignment between objective evidence and choice relative to control, whereas no comparable effect was observed for incongruent manipulation. Bars indicate medians, error bars show half the interquartile range, and dots represent individual participants. Asterisks denote significant differences (Wilcoxon signed-rank test, p<0.05). (**i,j**) Relationship between each participant’s saliency-congruency index and discriminability between the two most informative cues (S1′ and S4′), measured using learned cue values (i) or participants’ reported posterior estimates (j). Solid red lines indicate regression fits with shaded 95% confidence intervals; gray lines indicate baseline discriminability in the absence of saliency manipulation. No significant relationship was observed.

**Figure S10.**
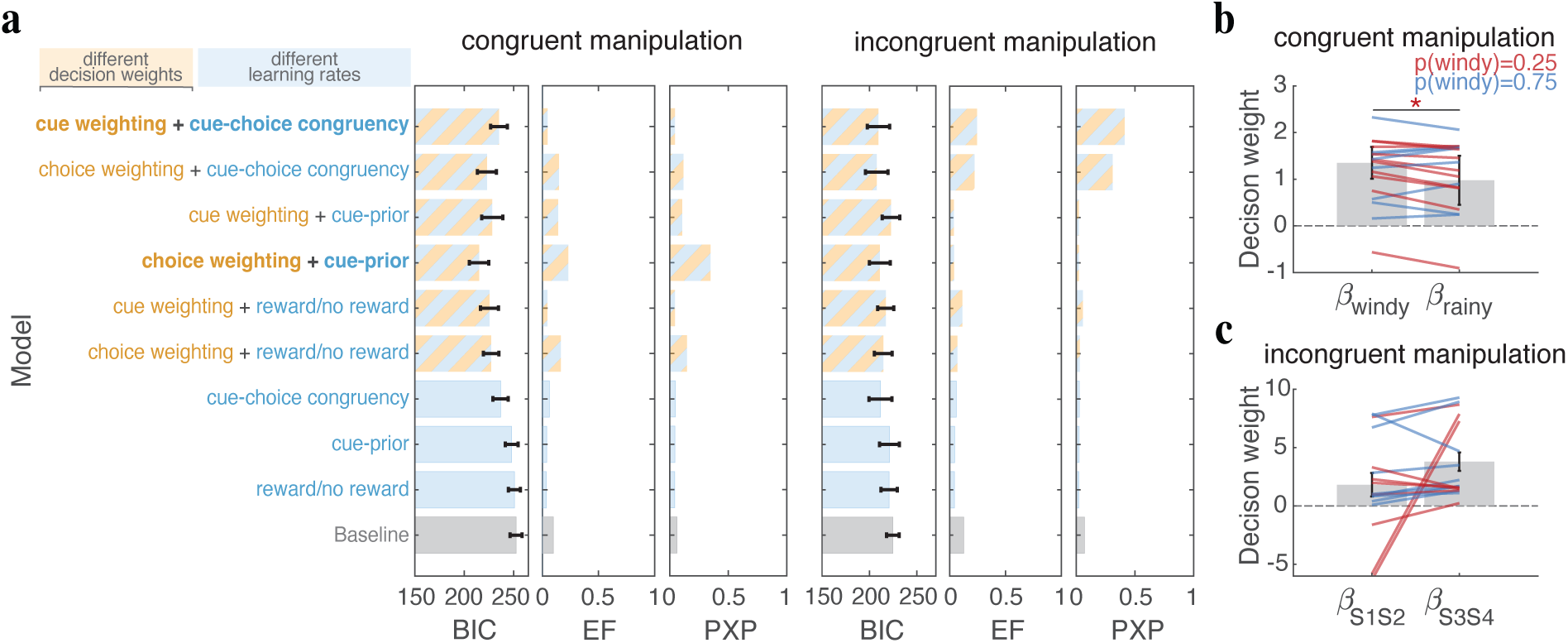
Distinct computational models account for behavior under congruent and incongruent saliency manipulations. (**a**) Model comparison for the congruent and incongruent manipulation groups using BIC, estimated frequency (EF), and protected exceedance probability (PXP). Lower BIC values indicate better fit, whereas higher EF and PXP indicate stronger evidence for a model. Participants in the congruent manipulation group were best fit by the cue-prior model with differential cue weighting during decision making, whereas participants in the incongruent manipulation group were best fit by models incorporating cue-choice congruency-dependent learning with either cue-or option-specific decision weighting. (**b**) Estimated decision weights from the winning model for the congruent manipulation group. (**c**) Estimated decision weights from the winning model for the incongruent manipulation group.

**Table S1.** Predicting choice from dwell time during the choice epoch. Trial-by-trial logistic regression models were used to predict participants’ choices from differential gaze toward the cue groups, the choice options, or both. Predictive accuracy was evaluated using within-subject 5-fold cross-validation. Reported t-and d-values reflect comparisons of each model against the null model. Gaze toward the choice options significantly improved prediction accuracy relative to the null model, as did the combined model. In contrast, gaze toward the cues alone did not improve prediction accuracy beyond the null model, and adding cue gaze to option gaze did not improve prediction beyond the choice-options model alone (t(72)=0.44, p=0.664, d=0.05). At the group level, however, mixed-effects logistic regression showed that both differential gaze toward choice options (b=2.79, SE=0.07, p<.001) and differential gaze toward cue groups (b=0.46, SE=0.05, p<.001) were significantly associated with choice when included together in the combined model. These results indicate that although gaze toward both cues and options relates to choice behavior, gaze toward cues does not explain additional variance beyond what participants learned during the task, whereas gaze toward the choice options provides additional predictive information about choice.

| *Model* | *Accuracy (%)* | *Statistic* | *p* | *d* |
| --- | --- | --- | --- | --- |
| Null | 70.95 | — | — | — |
| Cues | 70.87 | $t(72) = -0.45$ | 0.648 | -0.05 |
| Choice options | 74.90 | $t(72) = 5.01$ | $3.73 \times 10^{-6}$ | 0.58 |
| Combined | 75.04 | $t(72) = 4.97$ | $4.26 \times 10^{-6}$ | 0.58 |

**Table S2.** Effect of cue–choice congruency (CC) asymmetry on task performance. Results of generalized linear models examining the relationship between CC asymmetry group and task performance during the early (first 70 trials) and late (final 70 trials) phases of learning. CC group was entered as a categorical predictor. During the early phase, performance differed significantly across CC groups, with the high-CC group outperforming both the low-CC and mid-CC groups. During the late phase, the overall model was not significant, although the high-CC group continued to outperform the low-CC group, whereas the difference between the high-and mid-CC groups was no longer significant.

| *Predictor* | *b* | *SE* | *Statistic* | *p* |
| --- | --- | --- | --- | --- |
| **Early phase** |  |  |  |  |
| overall model | | | F(2,70) = 14 | $7.6 \times 10^{-6}$ |
| Intercept | 0.801 | 0.019 | t(70) = 41.06 | $1.01 \times 10^{-50}$ |
| Low CC | -0.145 | 0.027 | t(70) = -5.25 | $1.49 \times 10^{-6}$ |
| Mid CC | -0.058 | 0.027 | t(70) = -2.13 | 0.036 |
| **Late phase** |  |  |  |  |
| overall model |  |  | F(2,70) = 2.87 | 0.063 |
| Intercept | 0.864 | 0.016 | t(70) = 52.54 | $5.55 \times 10^{-58}$ |
| Low CC | -0.054 | 0.023 | t(70) = -2.35 | 0.021 |
| Mid CC | -0.018 | 0.023 | t(70) = -0.81 | 0.422 |

**Table S3.** Effect of cue–choice congruency (CC) asymmetry on sensitivity to evidence. Results of generalized linear models (GLMs) examining the relationship between CC asymmetry group and sensitivity to evidence during the early (first 70 trials) and late (final 70 trials) phases of learning. Sensitivity to evidence was quantified as the Spearman rank correlation between the summed objective weight of evidence (sum logLR) carried by the presented cues and participants’ choices. CC group was entered as a categorical predictor. During the early phase, sensitivity to evidence differed significantly across CC groups, with the high-CC group showing greater sensitivity than both the low-CC and mid-CC groups. During the late phase, the same directional pattern remained, with participants exhibiting stronger CC effects tending to make more evidence-consistent choices; however, the overall model was not significant, and neither the low-CC nor mid-CC group differed significantly from the high-CC group.

| *Predictor* | *b* | *SE* | *Statistic* | *p* |
| --- | --- | --- | --- | --- |
| **Early phase** |  |  |  |  |
| overall model |  |  | F(2,70) = 22.1 | 3.54 × 10<sup>-8</sup> |
| Intercept | 0.551 | 0.032 | t(70) = 17.10 | 1.07 × 10<sup>-26</sup> |
| Low CC | -0.303 | 0.045 | t(70) = -6.654 | 5.29 × 10<sup>-9</sup> |
| Mid CC | -0.148 | 0.045 | t(70) = -3.297 | 0.001 |
| **Late phase** |  |  |  |  |
| overall model |  |  | F(2,70) = 2.7 | 0.074 |
| Intercept | 0.554 | 0.037 | t(70) = 14.88 | 2.25 × 10<sup>-23</sup> |
| Low CC | -0.099 | 0.053 | t(70) = -1.89 | 0.063 |
| Mid CC | -0.011 | 0.052 | t(70) = 0.21 | 0.833 |

## Supplementary Note 1: Dwell time reflects learned evidence and decision-making processing

Because visual fixation enhances neural processing of attended stimuli (Desimone & Duncan, 1995; J. H. Reynolds & Chelazzi, 2004), and choice-related processes guide overt attention (Anderson, 2013; Cavanagh et al., 2014; Krajbich et al., 2010), gaze behavior can provide an indirect measure of which cue-and choice-related representations are preferentially processed during different phases of a trial. In particular, coordinated activation of cue-and choice-related representations may be reflected in patterns of dwell time. To examine this, we quantified the proportion of dwell time allocated to each cue and choice option relative to the total dwell time on all task-relevant elements within each trial epoch, averaged across trials.

We first asked whether participants preferentially allocated dwell time to cues supporting the higher-prior option. During the feedback epoch, participants showed a modest tendency to spend more time on cues favoring the higher-prior option, whereas no such bias was observed during the choice epoch (**Fig. S5a,d**). Specifically, participants with p(windy)=0.75 spent more time during feedback on S1 and S2 than on S3 and S4 (z=2.10, p=0.035, d=0.34), whereas no difference was observed during the choice epoch (z=0.12, p=0.898, d=0.02). Participants with p(windy)=0.25 showed the same directional trend during feedback, spending more time on S3 and S4 than on S1 and S2, although this effect did not reach significance (z=1.14, p=0.251, d=0.19); again, no difference was observed during the choice epoch (z=−1.05, p=0.292, d=−0.17).

We also tested whether gaze was preferentially allocated to cues based on their overall informativeness, independent of the option they supported. Specifically, we compared dwell time between cues carrying stronger evidence (S1 and S4) and weaker evidence (S2 and S3). No consistent differences were observed in either the choice or feedback epochs (choice: p(windy)=0.25: z=−1.31, p=0.186, d=−0.22; p(windy)=0.75: z=0.59, p=0.551, d=0.09; feedback: p(windy)=0.25: z=−0.26, p=0.78, d=−0.04; p(windy)=0.75: z=−0.52, p=0.602, d=−0.08). Together, these findings suggest that dwell time during feedback is more closely related to the relevance of cues to the chosen or higher-prior option than to their absolute informational strength.

With respect to the choice options themselves, participants preferentially allocated dwell time to the higher-prior option during both the choice and feedback epochs, although this bias was substantially stronger during the choice epoch (choice: p(windy)=0.25: z=5.18, p=2.16×10^−7^, d=0.86; p(windy)=0.75: z=4.83, p=1.33×10^−6^, d=0.79; feedback: p(windy)=0.25: z=2.81, p=0.005, d=0.46; p(windy)=0.75: z=2.65, p=0.008, d=0.43; **Fig. S5b,e**).

Finally, we examined dwell time as a function of the relationship between the presented cues and the participant’s choice on each trial. Participants spent more time fixating cues that supported the chosen option (S1 and S2 when windy was chosen; S3 and S4 when rainy was chosen) than cues opposing it. This effect was observed during both the choice epoch (S1+S2: z=2.91, p=0.003, d=0.34; S3+S4: z=2.27, p=0.022, d=0.26; **Fig. S5c**) and the feedback epoch (S1+S2: z=2.26, p=0.023, d=0.26; S3+S4: z=2.25, p=0.024, d=0.26; **Fig. S5f**). Thus, gaze allocation showed a consistent bias toward cues congruent with the chosen option during both epochs of the experiment. These results suggest that gaze patterns (dwell time) reflect both learned evidence and choice-related processing, potentially supporting the coactivation of cue-and choice-related representations required for reward-dependent Hebbian plasticity.

## References

Anderson, B. A. (2013). A value-driven mechanism of attentional selection. Journal of Vision, 13(3), 7. 10.1167/13.3.7

Arbel, Y., Feeley, E., & He, X. (2020). The Effect of Feedback on Attention Allocation in Category Learning: An Eye Tracking Study. Frontiers in Psychology, 11. 10.3389/fpsyg.2020.559334

Behrens, T. E. J., Woolrich, M. W., Walton, M. E., & Rushworth, M. F. S. (2007). Learning the value of information in an uncertain world. Nature Neuroscience 2007 10:9, 10(9), 1214–1221. 10.1038/nn1954

Bruckner, R., Heekeren, H. R., Nassar, M. R., & Carney, N. D. (2025). Understanding learning through uncertainty and bias. Communications Psychology 2025 3:1, 3(1), 24. 10.1038/s44271-025-00203-y

Brzosko, Z., Mierau, S. B., & Paulsen, O. (2019). Neuromodulation of Spike-Timing-Dependent Plasticity: Past, Present, and Future. Neuron, 103(4), 563–581. 10.1016/j.neuron.2019.05.041

Cavanagh, J. F., Wiecki, T. V., Kochar, A., & Frank, M. J. (2014). Eye tracking and pupillometry are indicators of dissociable latent decision processes. Journal of Experimental Psychology. General, 143(4), 1476–1488. 10.1037/a0035813

Ciranka, S., Linde-Domingo, J., Padezhki, I., Wicharz, C., Wu, C. M., & Spitzer, B. (2022). Asymmetric reinforcement learning facilitates human inference of transitive relations. Nature Human Behaviour, 6(4), 555–564. 10.1038/s41562-021-01263-w

Collins, A. G. E., & Frank, M. J. (2012). How much of reinforcement learning is working memory, not reinforcement learning? A behavioral, computational, and neurogenetic analysis. The European Journal of Neuroscience, 35(7), 1024–1035. 10.1111/J.1460-9568.2011.07980.X

Desimone, R., & Duncan, J. (1995). Neural Mechanisms of Selective Visual Attention. Annual Review of Neuroscience, 18(Volume 18, 1995), 193–222. 10.1146/annurev.ne.18.030195.001205

Donahue, C. H., & Lee, D. (2015). Dynamic routing of task-relevant signals for decision making in dorsolateral prefrontal cortex. Nature Neuroscience 2015 18:2, 18(2), 295–301. 10.1038/nn.3918

Farashahi, S., Donahue, C. H., Hayden, B. Y., Lee, D., & Soltani, A. (2019). Flexible combination of reward information across primates. Nature Human Behaviour 2019 3:11, 3(11), 1215–1224. 10.1038/s41562-019-0714-3

Farashahi, S., Donahue, C. H., Khorsand, P., Seo, H., Lee, D., & Soltani, A. (2017). Metaplasticity as a Neural Substrate for Adaptive Learning and Choice under Uncertainty. Neuron, 94(2), 401–414.e6. 10.1016/j.neuron.2017.03.044

Farashahi, S., Rowe, K., Aslami, Z., Lee, D., & Soltani, A. (2017). Feature-based learning improves adaptability without compromising precision. Nature Communications 2017 8:1, 8(1), 1768. 10.1038/s41467-017-01874-w

Frémaux, N., & Gerstner, W. (2016). Neuromodulated Spike-Timing-Dependent Plasticity, and Theory of Three-Factor Learning Rules. Frontiers in Neural Circuits, 9. 10.3389/fncir.2015.00085

Gerstner, W., Lehmann, M., Liakoni, V., Corneil, D., & Brea, J. (2018). Eligibility Traces and Plasticity on Behavioral Time Scales: Experimental Support of NeoHebbian Three-Factor Learning Rules. Frontiers in Neural Circuits, 12. 10.3389/fncir.2018.00053

Iigaya, K. (2016). Adaptive learning and decision-making under uncertainty by metaplastic synapses guided by a surprise detection system. eLife, 5, e18073. 10.7554/eLife.18073

Izhikevich, E. M. (2007). Solving the Distal Reward Problem through Linkage of STDP and Dopamine Signaling. Cerebral Cortex, 17(10), 2443–2452. 10.1093/cercor/bhl152

Jang, A. I., Costa, V. D., Rudebeck, P. H., Chudasama, Y., Murray, E. A., & Averbeck, B. B. (2015). The Role of Frontal Cortical and Medial-Temporal Lobe Brain Areas in Learning a Bayesian Prior Belief on Reversals. The Journal of Neuroscience, 35(33), 11751– 11760. 10.1523/JNEUROSCI.1594-15.2015

Khorsand, P., & Soltani, A. (2017). Optimal structure of metaplasticity for adaptive learning. PLOS Computational Biology, 13(6), e1005630. 10.1371/journal.pcbi.1005630

Kim, S., Hwang, J., & Lee, D. (2008). Prefrontal Coding of Temporally Discounted Values during Inter-temporal Choice. Neuron, 59(1), 161–172. 10.1016/j.neuron.2008.05.010

Krajbich, I., Armel, C., & Rangel, A. (2010). Visual fixations and the computation and comparison of value in simple choice. Nature Neuroscience, 13(10), 1292–1298. 10.1038/nn.2635

Lefebvre, G., Summerfield, C., & Bogacz, R. (2022). A Normative Account of Confirmation Bias During Reinforcement Learning. Neural Computation, 34(2), 307–337. 10.1162/NECO_A_01455

Legenstein, R., Pecevski, D., & Maass, W. (2008). A Learning Theory for Reward-Modulated Spike-Timing-Dependent Plasticity with Application to Biofeedback. PLOS Computational Biology, 4(10), e1000180–e1000180. 10.1371/JOURNAL.PCBI.1000180

Leong, Y. C., Radulescu, A., Daniel, R., DeWoskin, V., & Niv, Y. (2017). Dynamic Interaction between Reinforcement Learning and Attention in Multidimensional Environments. Neuron, 93(2), 451–463. 10.1016/j.neuron.2016.12.040

McGinty, V. B. (2019). Overt Attention toward Appetitive Cues Enhances Their Subjective Value, Independent of Orbitofrontal Cortex Activity. eNeuro, 6(6). 10.1523/ENEURO.0230-19.2019

McGuire, J. T., Nassar, M. R., Gold, J. I., & Kable, J. W. (2014). Functionally dissociable influences on learning rate in a dynamic environment. Neuron, 84(4), 870–881. 10.1016/j.neuron.2014.10.013

Moran, R., Dayan, P., & Dolan, R. J. (2021). Human subjects exploit a cognitive map for credit assignment. Proceedings of the National Academy of Sciences of the United States of America, 118(4). 10.1073/PNAS.2016884118/-/DCSUPPLEMENTAL

Munet, N. T., & Wallis, J. D. (2026). Effects of overt and covert attention on decision-making dynamics in prefrontal cortex. bioRxiv, 2026.05.18.723036-2026.05.18.723036. 10.64898/2026.05.18.723036

Nassar, M. R., Rumsey, K. M., Wilson, R. C., Parikh, K., Heasly, B., & Gold, J. I. (2012). Rational regulation of learning dynamics by pupil-linked arousal systems. Nature Neuroscience 2012 15:7, 15(7), 1040–1046. 10.1038/nn.3130

Neftci, E. O., & Averbeck, B. B. (2019). Reinforcement learning in artificial and biological systems. Nature Machine Intelligence, 1(3), 133–143. 10.1038/s42256-019-0025-4

Niv, Y. (2019). Learning task-state representations. Nature Neuroscience, 22(10), 1544–1553. 10.1038/s41593-019-0470-8

Niv, Y., Daniel, R., Geana, A., Gershman, S. J., Leong, Y. C., Radulescu, A., & Wilson, R. C. (2015). Reinforcement learning in multidimensional environments relies on attention mechanisms. Journal of Neuroscience, 35(21), 8145–8157. Scopus. 10.1523/JNEUROSCI.2978-14.2015

Palminteri, S., & Lebreton, M. (2022). The computational roots of positivity and confirmation biases in reinforcement learning. Trends in Cognitive Sciences, 26(7), 607–621. 10.1016/j.tics.2022.04.005

Palminteri, S., Lefebvre, G., Kilford, E. J., & Blakemore, S.-J. (2017). Confirmation bias in human reinforcement learning: Evidence from counterfactual feedback processing. PLoS Computational Biology, 13(8), e1005684. 10.1371/journal.pcbi.1005684

Payzan-LeNestour, E., Dunne, S., Bossaerts, P., & O’Doherty, J. P. (2013). The Neural Representation of Unexpected Uncertainty during Value-Based Decision Making. Neuron, 79(1), 191–201. 10.1016/j.neuron.2013.04.037

Piray, P., & Daw, N. D. (2024). Computational processes of simultaneous learning of stochasticity and volatility in humans. Nature Communications, 15(1), 9073. 10.1038/s41467-024-53459-z

Pozzi, I., Bohte, S., & Roelfsema, P. (2020). Attention-Gated Brain Propagation: How the brain can implement reward-based error backpropagation. Advances in Neural Information Processing Systems, 33, 2516–2526. https://proceedings.neurips.cc/paper_files/paper/2020/hash/1abb1e1ea5f481b589da523 03b091cbb-Abstract.html

Reynolds, J. H., & Chelazzi, L. (2004). Attentional modulation of visual processing. Annual Review of Neuroscience, 27, 611–647. 10.1146/annurev.neuro.26.041002.131039

Reynolds, J. N. J., & Wickens, J. R. (2002). Dopamine-dependent plasticity of corticostriatal synapses. Neural Networks, 15(4–6), 507–521. 10.1016/S0893-6080(02)00045-X

Rigoux, L., Stephan, K. E., Friston, K. J., & Daunizeau, J. (2014). Bayesian model selection for group studies—Revisited. NeuroImage, 84, 971–985. 10.1016/j.neuroimage.2013.08.065

Roelfsema, P. R., & Ooyen, A. van. (2005). Attention-Gated Reinforcement Learning of Internal Representations for Classification. Neural Computation, 17(10), 2176–2214. 10.1162/0899766054615699

Rushworth, M. F. S., & Behrens, T. E. J. (2008). Choice, uncertainty and value in prefrontal and cingulate cortex. Nature Neuroscience, 11(4), 389–397. 10.1038/nn2066

Samejima, K., Ueda, Y., Doya, K., & Kimura, M. (2005). Representation of action-specific reward values in the striatum. Science (New York, N.Y.), 310(5752), 1337–1340. 10.1126/science.1115270

Schultz, W. (2013). Updating dopamine reward signals. Current Opinion in Neurobiology, 23(2), 229–238. 10.1016/j.conb.2012.11.012

Schultz, W., Dayan, P., & Montague, P. R. (1997). A neural substrate of prediction and reward. Science, 275(5306), 1593–1599. 10.1126/SCIENCE.275.5306.1593;JOURNAL:JOURNAL:SCIENCE;WGR OUP:STRING:PUBLICATION

Shindou, T., Shindou, M., Watanabe, S., & Wickens, J. (2019). A silent eligibility trace enables dopamine-dependent synaptic plasticity for reinforcement learning in the mouse striatum. European Journal of Neuroscience, 49(5), 726–736. 10.1111/ejn.13921

Shouval, H. Z., & Kirkwood, A. (2025). Eligibility traces as a synaptic substrate for learning. Current Opinion in Neurobiology, 91, 102978. 10.1016/j.conb.2025.102978

Simoens, J., Verguts, T., & Braem, S. (2024). Learning environment-specific learning rates. PLoS Computational Biology, 20(3). 10.1371/JOURNAL.PCBI.1011978

Smith, S. M., & Krajbich, I. (2019). Gaze Amplifies Value in Decision Making. Psychological Science, 30(1), 116–128. 10.1177/0956797618810521

Soltani, A., Khorsand, P., Guo, C., Farashahi, S., & Liu, J. (2016). Neural substrates of cognitive biases during probabilistic inference. Nature Communications, 7(1), 11393. 10.1038/ncomms11393

Soltani, A., & Koechlin, E. (2021). Computational models of adaptive behavior and prefrontal cortex. Neuropsychopharmacology 2021 47:1, 47(1), 58–71. 10.1038/s41386-021-01123-1

Soltani, A., Murray, J. D., Seo, H., & Lee, D. (2021). Timescales of cognition in the brain. Current Opinion in Behavioral Sciences, 41, 30–37. 10.1016/j.cobeha.2021.03.003

Soltani, A., & Wang, X. J. (2006). A Biophysically Based Neural Model of Matching Law Behavior: Melioration by Stochastic Synapses. Journal of Neuroscience, 26(14), 3731– 3744. 10.1523/JNEUROSCI.5159-05.2006

Soltani, A., & Wang, X. J. (2008). From biophysics to cognition: Reward-dependent adaptive choice behavior. Current Opinion in Neurobiology, 18(2), 209–216. 10.1016/J.CONB.2008.07.003

Soltani, A., & Wang, X.-J. (2010). Synaptic computation underlying probabilistic inference. Nature Neuroscience, 13(1), 112–119. 10.1038/nn.2450

Stephan, K. E., Penny, W. D., Daunizeau, J., Moran, R. J., & Friston, K. J. (2009). Bayesian model selection for group studies. NeuroImage, 46(4), 1004–1017. 10.1016/j.neuroimage.2009.03.025

Sutton, R. S., & Barto, A. G. (1998). Reinforcement Learning: An Introduction. MIT Press.

Vartak, D., Jeurissen, D., Self, M. W., & Roelfsema, P. R. (2017). The influence of attention and reward on the learning of stimulus-response associations. Scientific Reports, 7(1), 9036. 10.1038/s41598-017-08200-w

Wang, M. C., & Soltani, A. (2025). Contributions of Attention to Learning in Multidimensional Reward Environments. Journal of Neuroscience, 45(7). 10.1523/JNEUROSCI.2300-23.2024

Yazdanpanah, A., Jung, H., Soltani, A., & Wager, T. D. (2026). Social information creates self-fulfilling prophecies in judgments of pain, vicarious pain, and cognitive effort. Proceedings of the National Academy of Sciences of the United States of America, 123(7), e2513856123–e2513856123. 10.1073/PNAS.2513856123;WGROUP:STRING:PUBLICATION

Yoo, A. H., & Collins, A. G. E. (2022). How Working Memory and Reinforcement Learning Are Intertwined: A Cognitive, Neural, and Computational Perspective. Journal of Cognitive Neuroscience, 34(4), 551–568. 10.1162/JOCN_A_01808

